# Single-cell transcriptomic profiling maps monocyte/macrophage transitions after myocardial infarction in mice

**DOI:** 10.1101/2020.04.14.040451

**Authors:** Giuseppe Rizzo, Ehsan Vafadarnejad, Panagiota Arampatzi, Jean-Sébastien Silvestre, Alma Zernecke, Antoine-Emmanuel Saliba, Clément Cochain

## Abstract

**Rationale:** Monocytes and macrophages have a critical and dual role in post-ischemic cardiac repair, as they can foster both tissue healing and damage. To decipher how monocytes/macrophages acquire heterogeneous functional phenotypes in the ischemic myocardium, we profiled the gene expression dynamics at the single-cell level in circulating and cardiac monocytes/macrophages following experimental myocardial infarction (MI) in mice.

**Methods and results:** Using time-series single-cell transcriptome and cell surface epitope analysis of blood and cardiac monocytes/macrophages, as well as the integration of publicly available and independently generated single-cell RNA-seq data, we tracked the transitions in circulating and cardiac monocyte/macrophage states from homeostatic conditions up to 11 days after MI in mice. We show that MI induces marked and rapid transitions in the cardiac mononuclear phagocyte population, with almost complete disappearance of tissue resident macrophages 1 day after ischemia, and rapid infiltration of monocytes that locally acquire discrete and time-dependent transcriptional states within 3 to 7 days. Ischemic injury induced a shift of circulating monocytes towards granulocyte-like transcriptional features (*Chil3, Lcn2, Prtn3*). Trajectory inference analysis indicated that while conversion to Ly6C^low^ monocytes appears as the default fate of Ly6C^hi^ monocytes in the blood, infiltrated monocytes acquired diverse gene expression signatures in the injured heart, notably transitioning to two main MI-associated macrophage populations characterized by MHCII^hi^ and *Trem2*^*hi*^*Igf1*^*hi*^ gene expression signatures. Minor ischemia-associated macrophage populations with discrete gene expression signature suggesting specialized functions in e.g. iron handling or lipid metabolism were also observed. We further identified putative transcriptional regulators and new cell surface markers of cardiac monocyte/macrophage states.

**Conclusions:** Altogether, our work provides a comprehensive landscape of circulating and cardiac monocyte/macrophage states and their regulators after MI, and will help to further understand their contribution to post-myocardial infarction heart repair.

## Introduction

Macrophages are critically involved in post-myocardial infarction (MI) cardiac repair, where they have a dual role as they can either promote tissue repair or precipitate myocardial damage (1). In the acute post-MI phase, the dynamics and cellular mechanisms governing cardiac macrophage heterogeneity as well as acquisition of specific states harboring pro-healing or pathogenic functional capacities are not fully understood.

Recent studies have pinpointed macrophage ontogeny as a potential driver of cardiac macrophage phenotype, and indicate that tissue resident macrophages that self-renew independently of monocyte input have cardioprotective functions (2), (3), (4). After infarction, the abundance of resident macrophages in the ischemic area drastically drops, and macrophages in the infarcted myocardium mostly derive from recruited monocytes that massively infiltrate the heart in the first week after MI (2), (3), (5), (6). In contrast to tissue resident subsets, monocyte-derived macrophages are thought to promote inflammation and tissue damage in the murine or human heart (2),(3),(4),(7). However, several lines of evidence indicate that monocyte-derived macrophages not only foster tissue damage, but are also required for post-MI cardiac repair. Reduced monocyte supply to the heart (6) or defects in emergency monopoiesis (8) result in defective post-MI cardiac healing. Although they are initially disposed towards pro-inflammatory activities, cardiac infiltrating Ly6C^hi^ monocytes gradually differentiate into pro-tissue healing F4/80^hi^Ly6C^lo^ macrophages (9), and impairment in this differentiation process hampers myocardial repair (10),(11). Recent studies further indicate that tissue injury causes emergence of functionally distinct atypical monocyte subsets in the periphery such as segregated-nucleus-containing atypical monocytes (SatM) (12), Ym1^+^ monocytes (13), or monocytes displaying a granulocyte-like state (14), but whether this occurs after MI is unknown. Given the preponderance of monocyte derived macrophages in the post-MI heart and their dual role as actors of the healing process or perpetrators of tissue damage, it is crucial to precisely understand the mechanisms regulating monocyte-derived macrophage acquisition of specific states associated with pro-repair or pathogenic functional capacities.

Here, we investigated the local and systemic transcriptional landscape of monocyte/macrophage transitions and tissue specification in the acute phase after MI. Using time-series single-cell transcriptomic profiling with simultaneous cell surface epitope labeling (CITE-seq (15)), and integration of independently generated scRNA-seq datasets, we established a comprehensive census of cardiac monocyte/macrophage transitions and gene expression dynamics in the blood and heart following MI.

## Materials and methods

Detailed experimental methods are available with the Online Supplement.

All animal studies and numbers of animals used conform to the Directive 2010/63/EU of the European Parliament and have been approved by the appropriate local authorities (Regierung von Unterfranken, Würzburg, Germany, Akt -Z. 55.2-DMS-2532-2-743). Single-cell RNA-sequencing data generated for this report has been deposited in Gene Expression Omnibus (GSE135310) and will be made available upon publication. The datasets can browsed in a web-accessible interface:

**Figure 1:** https://infection-atlas.org/4099491356/

**Figure 2:** Monocytes/macrophages alone : https://infection-atlas.org/9074526738/

Monocytes/macrophages and neutrophils: https://infection-atlas.org/4629609923/

**Figure 5:** https://infection-atlas.org/6970782209/

## Results

### Analysis of time-dependent monocyte and macrophage heterogeneity in the ischemic heart

Using droplet-based scRNA-seq (10x Genomics platform), we first profiled 4,445 total cardiac CD45^+^ cells isolated at 0 (no MI), 1, 3, 5, and 7 days after MI in mice (**Table 1, Figure S1A**). We identified T cells (*Cd3d*), natural killer cells (*Nkg7*), B cells (*Cd79a, Cd79b*), dendritic cells (DCs) (*Cd209a, Itgax, Flt3*), monocytes/macrophages (*Lyz2, Fcgr1, Csf1r*), neutrophils (*S100a8*), and minute contaminations by non-leukocytes (*Igfbp7, Sparc, Kdr*) and erythrocytes (*Hba-a2*) (**Figure S1B-C**). Monocytes, macrophages and neutrophils were extracted and examined in a new clustering analysis (**Figure 1** and **S1C**). Time-dependent neutrophil clusters (cluster 9, 10 and 13, **Figure 1A-C**) are described in detail in (16). Here, we focused our analysis on monocytes and macrophages (2,562 cells) and their heterogeneity over the post-MI time continuum (**Figure 1A-C, Table S1**).

**Figure 1:**
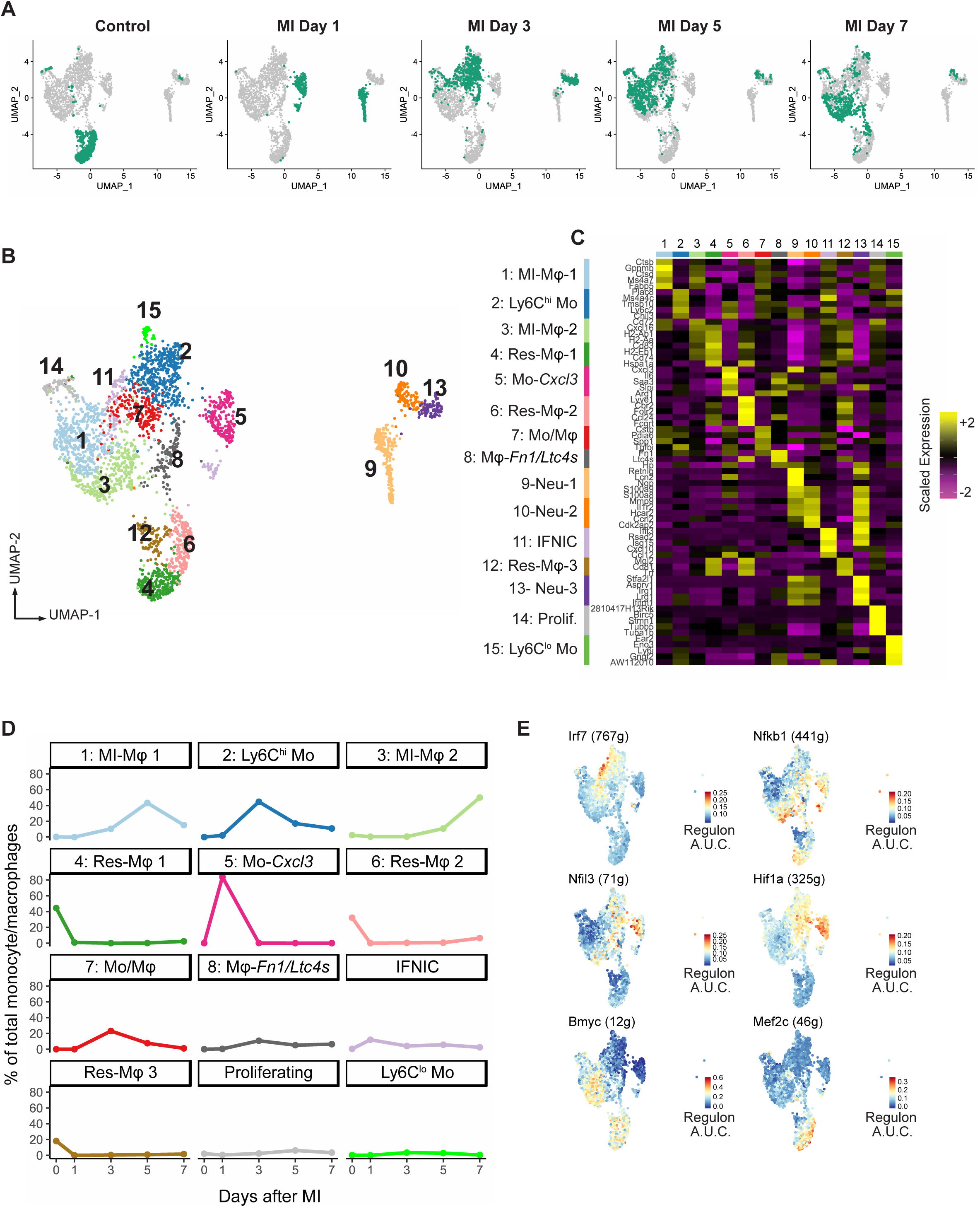
Time-resolved census of cardiac myeloid cells in the healthy and infarcted heart. UMAP of 3,012 monocytes/macrophages and neutrophils isolated from the healthy heart and at 1, 3, 5 and 7 days after MI with **A)** time point of origin and **B)** clustering analysis. **C)** Heatmap of the top 5 genes (ordered by log2 fold change) in each cluster; **D)** proportion of each cluster among total monocytes/macrophages according to time point after MI; **E)** regulon activity as predicted in SCENIC (A.U.C.: area under the curve) projected onto the UMAP plot.

**Table 1:** Murine myocardial infarction scRNA-seq datasets generated and analyzed. Mo/Mφ=monocyte/macrophages.*Included in analysis after quality control filtering **Median

| Reference | Organ | Data access | Technology | Time points | Sorting strategy | n cells * | Genes/cell in Mo/Mφ** | Sex |
| --- | --- | --- | --- | --- | --- | --- | --- | --- |
| New Data<br>(Figure 1) | Heart | GSE135310 | 10x Genomics Single Cell 3' v2 | Day 0, 1, 3, 5, 7 | Live CD45+ | Total: 4,445<br>Mo/Mφ: 2,562 | 2,200 | male |
| New Data<br>(Figure 2) | Heart | GSE135310 | 10x Genomics Single Cell 3' v3/CITE-seq/Cell Hashing | Day 1, 3, 5 | Live CD11b+ | Total: 3,402<br>Mo/Mφ: 1,904 | 4,685 | male |
| New Data<br>(Figure 5) | Blood<br>Heart | GSE135310 | 10x Genomics Single Cell 3' v3/CITE-seq/Cell Hashing | Control, Sham Day 1/3, MI day 1/3 | Live CD19-<br>NK1.1-Ter119-<br>CD11b+ | Total: 9,848<br>Mo/Mφ: 3,378 | 3,339 | male |
| Dick et al. Nat Immunol 2019 | Heart | GSE119355 | 10x Genomics Single Cell 3' v2 | Day 0, 11 | Live<br>CD45+CD64+C<br>D11b+ | Total: 5,802<br>Mo/Mφ: 5,802 | 2,504 | female |
| Farbehi et al. eLife 2019 | Heart | E-MTAB-7376 | 10x Genomics Single Cell 3' v2 | Day 0, 3, 7 | All live interstitial cells | Total: 14,041<br>Mo/Mφ: 4,004 | 1,846 | male |

Mo/Mφ=monocyte/macrophages
\*\*Median

Before MI, 3 resident macrophage (Mφ) clusters (clusters 4, 6 and 12, **Figure 1B-C**) were found. Res-Mφ1 expressed MHCII encoding genes (*Cd74, H2-Eb1*), while Res-Mφ2 expressed *Timd4* and *Lyve1* (**Figure 1B-D**), hence corresponding to TIMD4^neg^Lyve1^neg^MHCII^hi^ and TIMD4^+^Lyve1^+^MHCII^low^ tissue resident Mφ, respectively (2),(4). Res-Mφ3 displayed significant enrichment in few marker genes and may represent an intermediate resident Mφ population. Proportions of all these clusters dropped drastically after MI, consistent with previous observations (2) (**Figure 1D**).

At 1 and 3 days after MI, 2 monocyte populations (clusters 2 and 5, **Figure 1B-D**) were predominant and clearly differed in their gene expression profile according to time. At day 1, monocytes were enriched for *Cxcl3, Il6, Saa3* or *Arg1* (cluster 5: Mo-*Cxcl3*, **Figure 1B-D**), whereas classical Ly6C^hi^ monocytes (*Ly6c2, Plac8, Chil3, Ccr2*) were preponderant at day 3 (cluster 2: Ly6C^hi^ Mo, **Figure 1B-D**). In addition, Ly6C^lo^ monocytes were found at low levels at 3 and 5 days post-MI (cluster 15: Ly6C^lo^ Mo, **Figure 1B-D**). A cluster with gene expression suggestive of an intermediate monocyte to macrophage differentiation state was also observed (Cluster 7: MI-Mo/Mφ, **Figure 1B-D**).

In contrast to day 1 and 3, two major MI-associated macrophage clusters (*Adgre1, C1qa*) were preponderant at days 5 and 7 post-MI (**Figure 1A-D**). MI-Mφ-1 had higher expression of *Trem2, Cd63, Fabp5, Gpnmb* and some known regulators of cardiac repair such as *Timp2, Igf1* or *Spp1* (17), (18), (19) (**Figure 1C** and **S2**, “*Trem2/Igf1*^*hi*^” signature), while MI-Mφ-2 had higher expression of MHCII encoding genes (*Cd74, H2-Aa*) and pro-inflammatory genes (*Nlrp3, Il1b, Tlr2*), but also of some anti-inflammatory cytokines (*Il10*) (**Figure 1C** and **S2**, “MHCII” signature). Additional minor clusters included type I interferon response signature macrophages (IFNICs (20), cluster 11, *Isg15, Rsad2, Cxcl10*), and a macrophage cluster characterized by co-enrichment in *Fn1, Ltc4s* and *Arg1* (cluster 8, Mφ-*Fn1/Ltc4s*) (**Figure 1B-C**).

Single-cell regulatory network inference and clustering (SCENIC) (21) analysis revealed patterning of specific regulons activities (i.e. co-expression of transcription factors and their target genes) in IFNICs (e.g. *Irf7*), tissue resident macrophages (e.g. *Mef2c*), MI-associated macrophages (e.g. *Bmyc, Nfkb1*), and monocytes (e.g. *Hif1a, Nfil3*) (**Figure 1E**). Hence, our results indicate that shifts in monocyte/macrophage transcriptional states after MI may rely on the activity of specific transcriptional regulators.

### CITE-seq analysis of monocyte/macrophage transitions in the acute post-MI phase

To validate and refine our analysis of the monocyte to macrophage transition in the acute phase after MI, we performed CITE-seq (cellular indexing of transcriptomes and epitopes by sequencing (15)) analysis of cardiac CD11b^+^ cells at 1, 3, and 5 days after MI, where all cells were multiplexed in a single scRNA-seq library using a cell hashing approach to reduce inter-sample variability and batch effects (22) (**Figure 2A-B and S3**). Within CD11b^+^ cells, we identified Ly6G^+^*S100a8*^hi^ neutrophils and cDC2 (CD11c^+^MHCII^+^, *Flt3, Cd209a*) (**Figure S3A-C**). NK cells (*Nkg7*) were filtered out in pre-processing steps. Monocytes and macrophages were identified by expression of established surface markers and their encoding transcripts (CD64/*Fcgr1*, Ly6C/*Ly6c2*, CX3CR1/*Cx3cr1*, MSR1/*Msr1*) (**Figure S3A-B**), gated out and further analyzed separately (1,904 cells, **Figure 2** and **S4, Table S2**). Distinct clusters of monocyte/macrophages were observed in varying proportions at the different time points after MI (**Figure 2C-E** and **S4**), showing clear gene expression similarities to our first dataset (**Figure 2E**).

**Figure 2:**
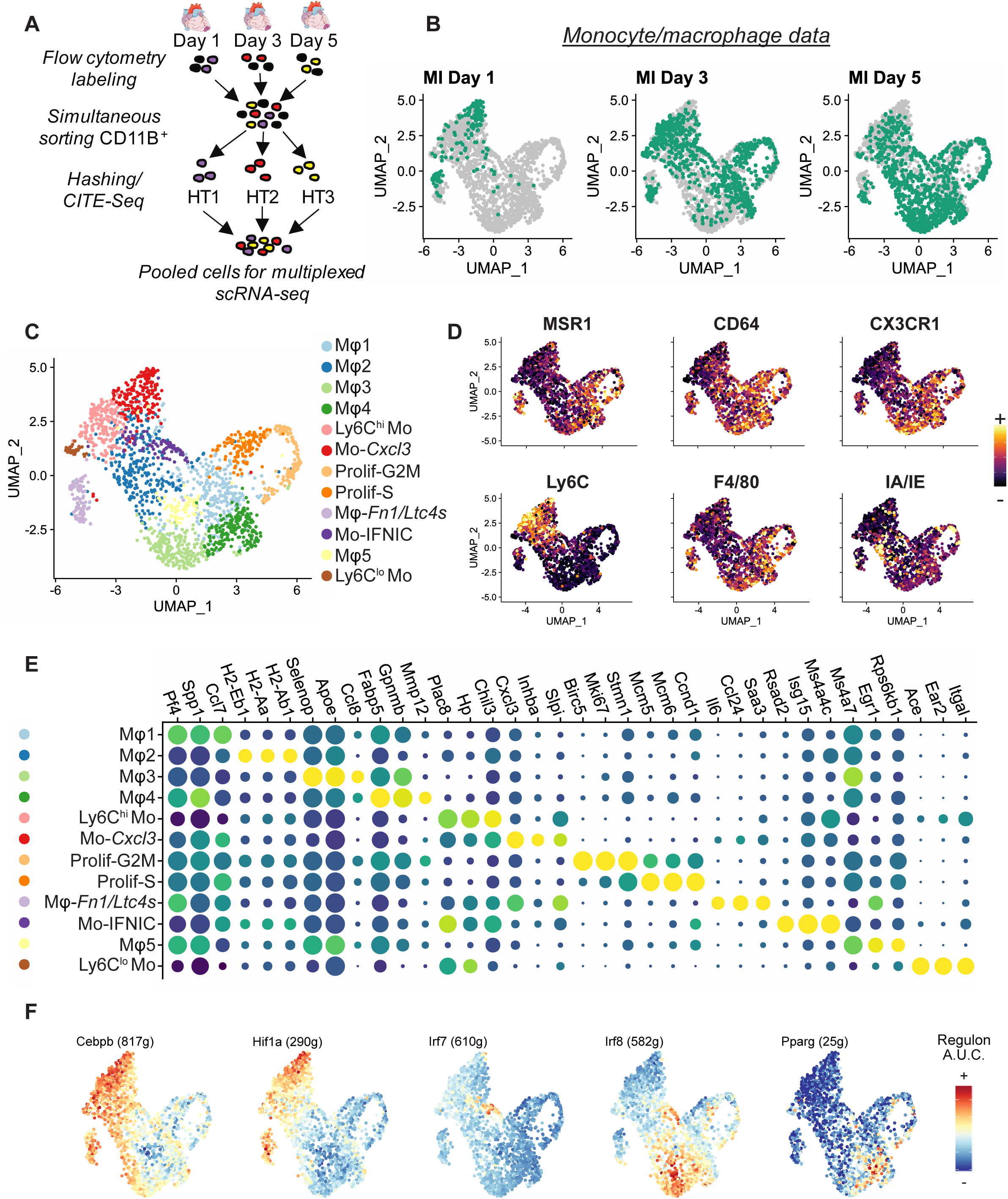
Multiplexed analysis of monocytes/macrophages at day 1, 3 and 5 after MI. **A)** Flowchart of the experimental design; **B)** time point of origin of single cells projected onto the UMAP plot; **C)** Seurat cluster annotation of monocytes/macrophages color-coded on the UMAP plot; **D)** expression of the indicated cell surface markers measured by CITE-seq projected onto the UMAP plot; **E)** dotplot showing expression of the top 3 enriched genes (ranked by Log2 Fold Change) in each cluster; **F)** regulon activity as predicted in SCENIC (A.U.C.: area under the curve) projected onto the UMAP plot (in brackets: number of genes in regulon).

At day 1, the major cell cluster was Ly6C^+^*Cxcl3*^*hi*^ monocytes (Mo-*Cxcl3*) (**Figure 2B-E and S4A-B**). We also recovered Mφ-*Fn1/Ltc4s*, Ly6C^hi^ monocytes, Ly6C^lo^ monocytes, and monocytes with the IFNIC signature (IFNIC-Mo). At 3 and 5 days after MI, five macrophage clusters were observed. The proportions of Mφ1 (*Pf4, Spp1*) peaked at day 3, likely representing a transition state. The proportion of Mφ2 (*Il1b*, MHCII encoding genes), Mφ3 (*Apoe, Ms4a7, Trem2, Igf1*), and Mφ4 (*Lipa, Gpnmb, Fabp5, Mmp12*) gradually increased until day 5. Mφ5 (*Egr1*) represented <10%of all cells at day 5. Clusters of proliferating cells were identified as S phase and G2M phase-proliferating macrophages by cell cycle scoring in Seurat (23) (**Figure 2C-E** and **S4C**). SCENIC revealed specific patterns and gradients of transcriptional regulators activity in monocyte/macrophages, with high activity of the *Cebpb* and *Hif1a* regulons in *Cxcl3*^*hi*^ monocytes and *Fn1/Ltc4s*^*hi*^ Mφ, *Irf7* in IFNIC-Mo, *Pparg, Irf8* and *Bhlhe41* in macrophages (**Figure 2F and S4D**). These data independently confirm the time-dependent heterogeneity of Ly6C^+^ monocytes in the very acute post-MI phase (1 to 3 days), the presence of minor populations of Ly6C^lo^ monocytes, IFNICs and Mφ*-Fn1/Ltc4s*^*hi*^ Mφ with highly specific gene signatures, and the major dichotomy in MI-associated macrophages (*Trem2*^*hi*^*Igf1*^*hi*^ vs MHCII signature).

### Integrated scRNA-seq analysis of the monocyte/macrophage landscape over the post-MI time continuum

To validate the conservation of monocyte/macrophage states across independent scRNA-seq studies, increase the number of analyzed cells, and extend our observations to later time points, we integrated our new data with published scRNA-seq datasets of cardiac macrophages from the steady state up to 11 days after infarction (2), (24) (**Table 1**). Separate analysis of these data recovered monocyte and macrophage populations consistent with our own observations (**Figure S5** and **supplementary text**). After curating these datasets to exclude granulocytes, DCs, non-leukocyte contamination and “hybrid cells” (i.e. cells enriched for markers of both Mφ and non-immune cells in the Farbehi et al. dataset (24)), we performed data integration in Seurat v3 (23), encompassing 14,272 monocytes/macrophages across 5 time points and 4 datasets (**Figure 3A-B**). Clustering analysis revealed 16 cell populations with time-specific distributions (**Figure 3A-D, Table S3**).

**Figure 3:**
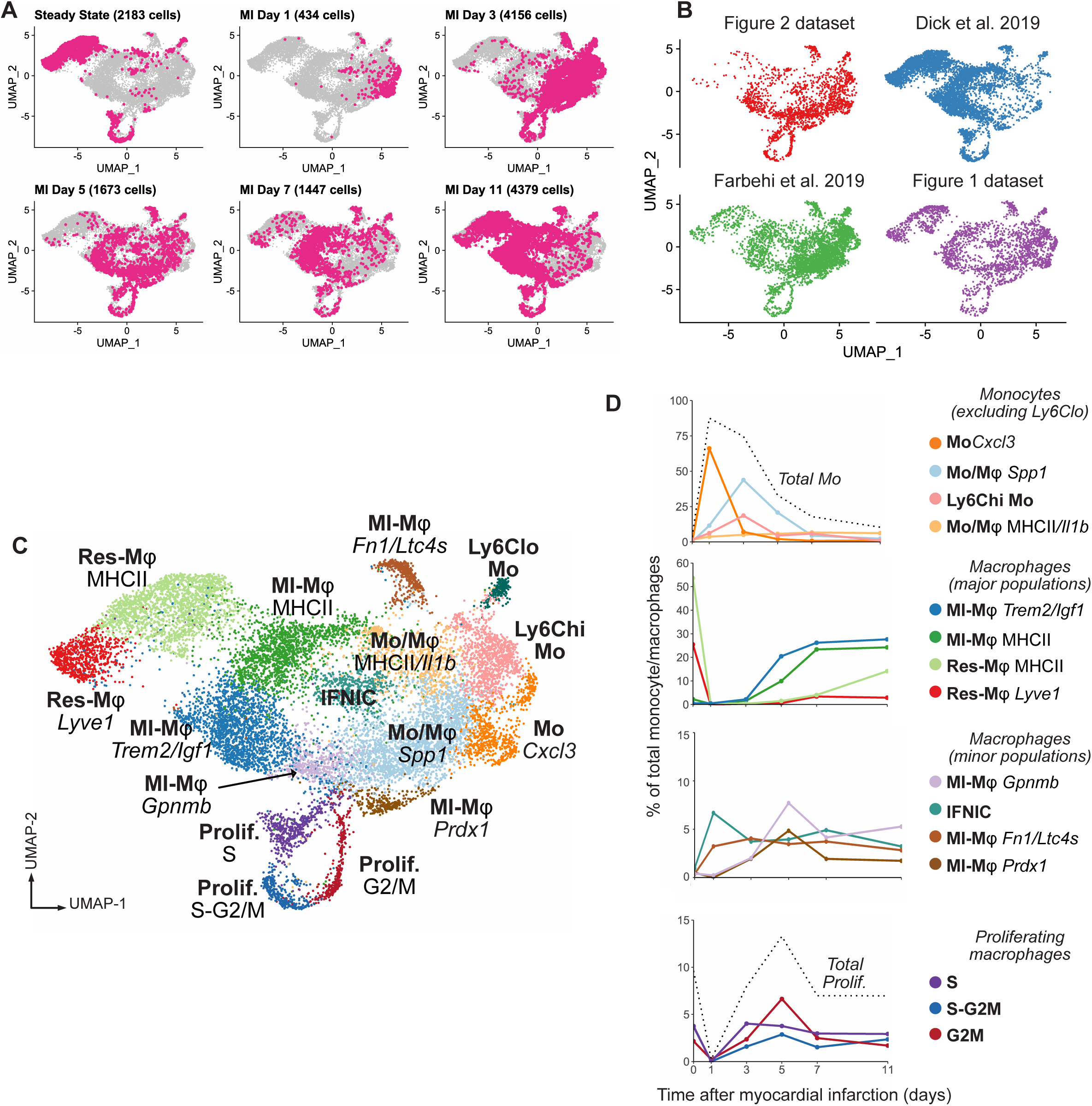
Monocyte/Macrophage transitions from the steady state to day 11 post-MI in 4 independent datasets. UMAP representation of single-cell RNA-seq gene expression data in 14,272 monocytes/macrophages from the healthy heart and at 1, 3, 5, 7 and 11 days after MI with **A)** time point of origin of each cell; **B)** dataset of origin of each cell and **C)** clustering analysis and identification of cell clusters. **D)** Proportion of the indicated clusters among total monocytes/macrophages according to time point after MI with a focus on monocytes (excluding Ly6C^low^ monocytes, dashed line= total Ly6C^+^ monocytes), major population of resident and MI-associated macrophages, minor populations of macrophages, and proliferating macrophage populations (dashed line= total proliferating macrophages).

We identified the two major tissue resident populations (Res-Mφ-*Timd4*, and Res-Mφ-MHCII), that were preponderant in the steady state heart. *Cxcl3* enriched monocytes (Mo-*Cxcl3*) peaked at 1 day. Two transitional monocyte/macrophage states were observed, characterized on one hand by enrichment for *Spp1*, and on the other hand for *Il1b* and MHCII encoding genes (**Figure 3C and S6**). At 5, 7 and 11 days after MI, two major MI-associated macrophage populations were observed (MI-Mφ-*Trem2/Ifg1*: *Apoe, Ms4a7*; and MI-Mφ-MHCII: *Cd74, H2-Aa*), consistent with our previous results (**Figure 3C-D, S6 and S7A**). Minor populations included MI-Mφ-*Prdx1* (also enriched for e.g. *Fth1, Ftl1, Slc48a1*), MI-Mφ-*Gpnmb* (*Fabp5, Lgals3, Lpl*), IFNICs and MI-Mφ-*Fn1/Ltc4s* (**Figure 3C** and **S6**). Although their represented under 10%of total monocyte/macrophages at all time points, all these populations increased in proportions after MI (**Figure 3D**). Gene ontology enrichment analysis (25), (26) (**Figure S8, Table S4**) revealed distribution of putative biological processes in the different populations, with e.g. MI-Mφ-*Prdx1* showing enrichment for functions related to cell detoxification and response to oxidative stress **(Figure S8**, blue frames), and molecular functions such as heme binding, iron binding, peroxiredoxin activity or glutathione transferase activity (**Figure S9, Table S5**).

Three clusters corresponded to proliferating macrophages and comprised a S phase population, an intermediate S/G2M population, and cells in the G2M phase (**Figure 3C** and **S7B**). Proliferating macrophage proportions drastically dropped at 1 day post-MI and rose thereafter to peak around day 5 (**Figure 3D**). Within proliferating macrophages, we identified cells corresponding to Res-Mφ-*Timd4* (*Lyve1, Timd4*) and Res-Mφ-MHCII (MHCII encoding genes, *Cd81, Mgl2, Lilra5*), IFNICs, MI-associated monocyte/macrophages enriched for MHCII encoding genes, *Il1b* and *Ccr2*, MI-associated monocyte/macrophages enriched for *Spp1, Hmox1, Cd63, Prdx1, Trem2* or *Igf1* abundant at day 3 and 5 after MI (**Figure S10**). These results indicate that cell proliferation feeds into most resident and MI-associated macrophage populations.

Compared to MI-associated monocytes and macrophages, tissue resident populations (Res-Mφ-*Timd4*, Res-Mφ-MHCII) shared enrichment for *Mgl2* (encoding CD301b) and *Cd81* **(Figure 4A)**. We evaluated whether these newly identified markers could be employed to track monocyte and macrophage transitions within the ischemic heart. In the steady state heart, flow cytometry revealed 3 major macrophages populations, characterized as MHCII^+^TIMD4^-^ (36.7%), MHCII^+^TIMD4^+^ (26.6%) and MHCII^-^TIMD4^+^ (25.3%) **(Figure 4B-E)**, Indicating that some intermediate states (i.e. the MHCII^+^TIMD4^+^ double positive population) may not generate transcriptionally distinct cell clusters in scRNA-seq analysis. Levels of these 3 populations dropped at 3 days post-MI, when most macrophages were MHCII^-^TIMD4^-^ **(Figure 4B-E)**. While tissue resident populations were mostly positive for surface MGL2 and CD81, with TIMD4^+^ populations showing the highest levels (**Figure 4F-G**), double negative MHCII^-^TIMD4^-^ lacked surface MGL2 and CD81 at day 3 (**Figure 4F-G**). Ly6C^hi^ monocytes were negative for MGL2 and CD81 (**Figure S7E**). At day 10 post-MI, macrophage counts were still slightly elevated, and consisted mostly of MHCII^-^TIMD4^-^ and MHCII^+^TIMD4^-^ cells with lower surface MGL2 and CD81 than their counterparts from control hearts (**Figure 4C-G**). Partial recovery of the MHCII^+^TIMD4^+^ and MHCII^-^TIMD4^+^ cells was observed (**Figure 4C-G**). This flow cytometry analysis confirms that MI induces a shift in macrophage populations lasting beyond the acute inflammatory phase, and uncovers the combination of MGL2 and CD81 surface expression as a potential new labeling strategy to discern tissue resident from recruited monocyte-derived macrophages in the ischemic heart.

**Figure 4:**
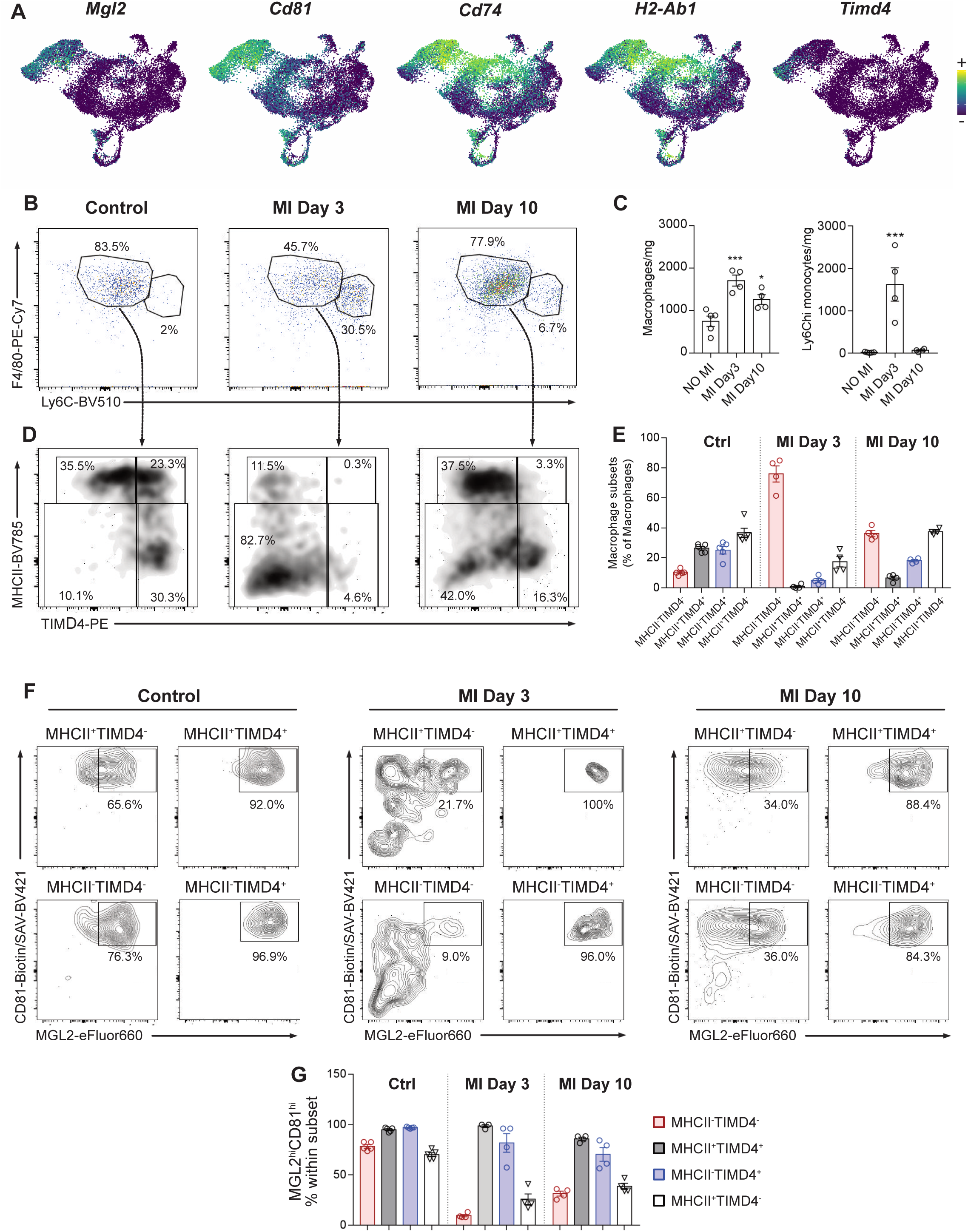
Flow cytometry analysis of monocyte and macrophage transitions. A) Expression of the indicated transcripts projected onto the UMAP plot of the integrated data (see **Figure 3**). **B)** Identification of F4/80^hi^Ly6C^lo^ macrophages and F4/80^low^Ly6C^hi^ monocytes (pre-gated on live CD45^+^CD11b^+^Ly6G^-^) and **C)** absolute counts in cells per mg in heart cell preparations in control hearts and at 3 and 10 days after MI (^*^p<0.05; ^***^p<0.001 versus No MI condition); **D)** identification of macrophage subsets based on MHCII and TIMD4 surface expression and **E)** proportion of these subsets within total macrophages; **F)** surface expression flow cytometry plots and **G)** proportion of MGL2^hi^CD81^hi^ cells in MHCII^-^TIMD4^-^, MHCII^+^TIMD4^+^, MHCII^-^TIMD4^+^ and MHCII^+^TIMD4^-^ macrophages at the indicated time points after MI.

### MI-Mφ-Trem2/Ifg1 and MI-Mφ-MHCII are of monocytic origin

The transcriptomic profile and time-dependent accumulation pattern of the two main MI-associated macrophage populations, MI-Mφ-MHCII and MI-Mφ-*Trem2/Ifg1*, clearly suggested that they originate from recruited monocytes. MI-Mφ-MHCII expressed the prototypical monocyte transcript *Ccr2* and although they had a gene expression profile similar to Res-Mφ-MHCII, they also expressed low levels of tissue-resident Mφ-specific transcripts such as *Mgl2, Cd81, Slco2b1* or *Lilra5* (**Figure S6-7**). MI-Mφ-*Trem2/Ifg1* had low *Ccr2* expression, but clearly differed from all resident cells as they had low levels of MHCII encoding transcripts, and of transcripts associated with Res-Mφ-*Timd4* (*Timd4, Lyve1, Cbr2, Fcgrt, Mgl2, Cd81, Slco2b1*) (**Figure S6-7**). TdTomato transcripts originating from *Cx3cr1* based lineage tracing of tissue resident macrophages in Dick et al. (2) mapped to Res-Mφ-*Timd4*, Res-Mφ-MHCII and proliferating macrophages, while only scattered cells were observed in MI-Mφ-*Trem2/Ifg1* and MI-Mφ-MHCII (**Figure S7C**). Although TdTomato transcript coverage was weak even in tissue resident macrophages (Res-Mφ-*Lyve1*=12.1%; Res-Mφ-MHCII: 9.2%; only cells from Dick et al. considered (2)) it was clearly much lower in MI-Mφ-*Trem2/Ifg1* (1.4%) and MI-Mφ-MHCII (1.0%) (2). These data indicate that MI-Mφ-*Trem2/Ifg1* and MI-Mφ-MHCII originate from recruited monocytes.

Altogether, our data provide a comprehensive single-cell map of cardiac monocyte/macrophage transitions over the post-MI time continuum, and demonstrate an ischemia induced shift in cardiac macrophage populations. An immediate reduction in tissue resident macrophage proportions was paralleled by Ly6C^+^ monocyte recruitment, with a striking time-dependent transcriptional heterogeneity of Ly6C^+^ monocytes in the first 3 days after MI. Subsequently, MI-associated macrophages originating from differentiated monocytes accumulated in the ischemic heart. We further identified minor MI-associated macrophage populations (IFNICs, Mφ-*Fn1/Ltc4s*, MI-Mφ-*Prdx1*), and characterized the main MI-associated macrophages as being defined by a MHCII^hi^ and a *Trem2*/*Igf1*^hi^ gene expression signature.

### CITE-seq analysis of circulating monocyte states after myocardial infarction

We then investigated whether the cardiac monocyte/macrophage states could be acquired remotely, notably via production of injury-associated atypical monocytes in the periphery that would transit in the bloodstream before infiltrating the myocardium. We performed CITE-seq analysis of CD19^-^NK1.1^-^Ter119^-^CD11b^+^ cells from the blood of naive control mice, sham-operated animals at 1 and 3 days post-surgery, and mice with MI at 1 and 3 days. We also included cardiac cells from the ischemic heart at 1 and 3 days post-MI. Samples were simultaneously analyzed using a cell hashing strategy (22) (**Figure 5A**).

**Figure 5:**
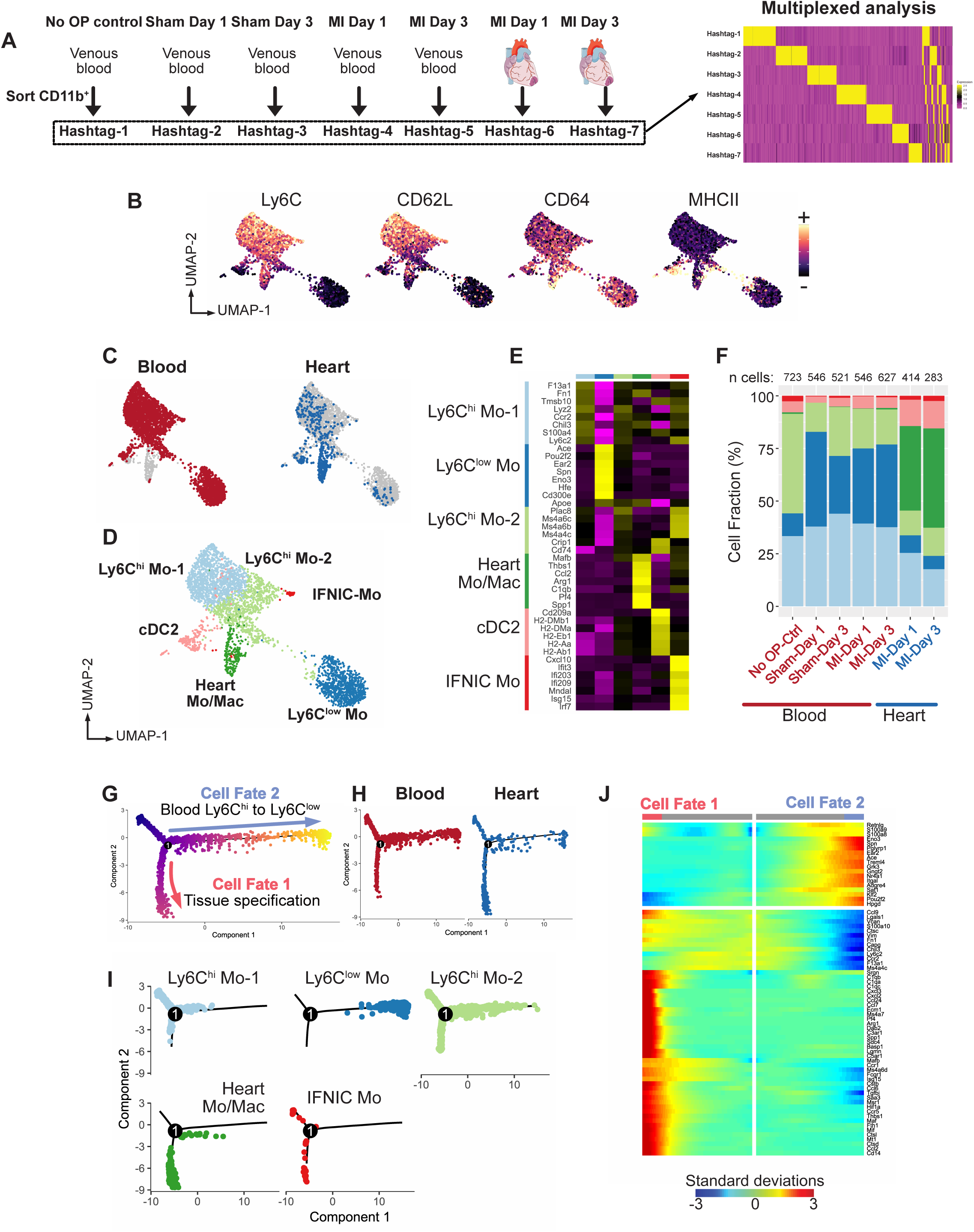
Single-cell RNA-seq analysis of the blood to heart transition in monocyte/macrophages. **A)** Experimental design overview; **B)** CITE-seq signal for the indicated surface markers in blood and heart monocyte/macrophages projected on the UMAP plot; **C)** tissue of origin projected on the UMAP plot; **D)** clustering analysis and identification of cell clusters; **E)** heatmap of average expression of the top 8 marker transcripts (ordered by fold change) per cluster (overlapping markers are shown only once); **F)** proportion of the indicated clusters among total monocytes/macrophages according to sample of origin; **G)** pseudotime analysis of monocytes/macrophages in Monocle, split according to **H)** tissue origin and **I)** Seurat clusters; **J)** heatmap of pseudotime gene expression variation on branches of the pseudotime tree (as indicated on panel G, only genes with q val <10^−150^ are shown).

Neutrophils were identified as CD115^neg^Ly6G^hi^, and minor contamination by T cells/NK cells, basophils/mast cells and B cells were detected (**Figure S11A-B**). We identified CD115^+^Ly6G^-^ monocytes (3,378 cells) that could further be divided into Ly6C^hi^ (2 clusters, also expressing CD62L) and Ly6C^low^ monocytes (1 cluster), MHCII^+^*Flt3*^+^*Cd209a*^+^*Clec10a*^+^ cDC2 (**Figure 5B-D, Figure S11A, Table S6**), and a minor population of CD115^+^Ly6C^hi^*Isg15*^hi^ Type I IFN response monocytes (IFNIC-Mo, **Figure 5C-E**). These populations were also found in infarct tissue samples, consistent with our previous results (**Figure 5C-F**).

One cell cluster contained almost only cells extracted from the ischemic heart (Heart Mo/Mac, **Figure 5C-F**). The 2 clusters of CD115^+^Ly6C^hi^ monocytes appeared to represent a continuum of gene expression states rather than truly distinct populations: Ly6C^hi^_Mo1 had higher expression of *Chil3* (encoding Ym1), and higher surface levels of Ly6C and CD62L (**Figure 5B and E, Table S7**), suggesting recent egress from the BM, while Ly6C^hi^_Mo2 were slightly enriched for marker genes of Ly6C^lo^ monocytes (*Treml4, Ace*) suggesting that they may represent an intermediate state of Ly6C^hi^ to Ly6C^lo^ monocyte conversion. Pseudotime analysis supported this notion, with a clear continuum from state Ly6C^hi^_Mo1 to Ly6C^lo^ monocytes, via Ly6C^hi^_Mo2, in the blood (**Figure 5G-J**). Along the Ly6C^hi^ to Ly6C^low^ blood differentiation axis (**Figure 5G-I**), cells acquired expression of e.g. *Cd36, Apoe, Nr4a1* or *Itgax* (**Figure 5I** and **S11D**), as previously described (27).

No clear injury-associated (i.e. caused by sham surgery) or MI-associated monocyte cluster was identified. However, within Ly6C^hi^ monocytes, MI (and sham surgery at day 1, likely reflecting transient systemic inflammation caused by thoracotomy (28)) appeared to cause a shift from Ly6C^hi^_Mo2 to Ly6C^hi^_Mo1, (**Figure 5F and S12A**), consistent with mobilization of recently produced monocytes from hematopoietic reservoirs after MI (6). Monocytes can arise from two independent differentiation pathways, i.e. monocyte-dendritic cell progenitors (MDPs) giving rise to mature monocytes with a “dendritic-cell like” profile, and granulocyte-monocyte progenitors (GMPs) giving rise to monocytes with a “granulocyte-like” state (14), (29). We detected expression of various granulocyte-associated transcripts such as *S100a8, S100a9, Lcn2* or *Mmp8* in the various monocyte populations, albeit at much lower levels than in neutrophils (**Figure S12B**). Based on the combined expression of 12 neutrophil characteristic transcripts, we applied a “Granulocyte Expression Score” to monocytes, which appeared highest in the Ly6C^hi^_Mo1 cluster. Within the Ly6C^hi^_Mo1 cluster, we observed a clear elevation of the Granulocyte Expression Score in cells from the sham and MI conditions at 1 and 3 days post-MI, with cells from the MI condition showing significantly higher Granulocyte Expression Score than their sham counterparts at both time point (**Figure S12C**). To avoid bias caused by surgery-induced transient inflammation, we analyzed differential gene expression in blood Ly6C^hi^_Mo1 from animals at day 3 after sham surgery or MI. Significantly upregulated genes in the MI condition included granulocyte-associated transcripts such as *Prtn3, Lrg1, Wfdc21* or *Lcn2*, as well as other transcripts potentially involved in inflammatory responses such as *Socs3* (**Figure S12D**). CD115^+^CD11b^+^Ly6C^hi^ mature monocytes sorted from the bone marrow of infarcted mice (day 3) had increased levels of *Prtn3, Lcn2* or *Chil3* **(Figure S12E**). Altogether, these results indicate that MI may shift monopoiesis towards the granulocyte-like differentiation pathway.

### Myocardial infarction does not induce appearance of segregated-nucleus-containing atypical monocytes (SatM) or type-I interferon signature monocytes in the circulation

Atypical fibrosis-associated SatM have been described and defined as F4/80^-^Ly6C^-^ Ceacam1^+^Msr1^+^ (12). Expression of *Ceacam1* was enriched in Ly6C^lo^ monocytes, which we confirmed at the protein level by flow cytometry in steady state blood monocytes (**Figure S12F**), but no circulating monocyte clearly corresponding to SatM could be observed in our single-cell analysis. A recent report proposed that ischemic injury induces type I IFN signaling that remotely primes a specific interferon-induced state in bone marrow progenitors, leading to appearance of monocytes with the IFNIC signature (30). In our analysis, IFNIC-Mo were already found in the steady state blood (2.62%of circulating monocytes/cDC2), and their proportions tended to decrease after MI (0.31 and 0.72%at day 1 and 3 post-MI) (**Figure 5F**). Also when analyzing data from the steady state condition alone, we found a discrete subset of CD115^+^Ly6C^+^ cells with a strong type I IFN signature (**Figure S13A-C**). We further assigned an ISG (interferon stimulated genes) score (30) to all cells, which as expected was highest in IFNIC-Mo (**Figure S13D**). In other monocyte populations, this score did not vary across experimental conditions (**Figure S13E**). We also detected a small cluster (cluster 12) with high expression of several ISGs (*ISG15, MX1, IFIT1, IFIT3, IFI6, IFI44*) in cells corresponding to CD3^-^CD19^-^CD56^-^CD14^+^CD16^-^ classical monocytes in CITE-seq analysis of peripheral blood mononuclear cells from a healthy human donor (**Figure S13F-H**; data from 10x Genomics). Altogether, these results indicate that MI does not induce the appearance of circulating SatM or IFNIC-type monocytes.

### Monocytes follow a tissue specification trajectory in the ischemic heart

We only recovered a low number of heart specific monocytes/macrophages in this experiment (264 cells at day 1, 283 cells at day 3). Nevertheless, we detected enrichment for markers of local tissue specification previously identified in cardiac monocyte/macrophage populations such as *Arg1, C1qb, Pf4, Cxcl3, Saa3* or *Spp1*, and increased surface levels of CD64 (**Figure 5B-E**). In trajectory inference analysis, these cells preferentially occupied a discrete branch of the pseudotime tree, suggesting a specific path of tissue specification of Ly6C^hi^ monocytes (**Figure 5G-J**).

Altogether, these data clarify the blood to heart transitions of myocardium infiltrating CD11b^+^ cells and show that (i) cDC2 and Ly6C^low^ monocytes that can be found in the heart in low numbers circulate in the blood already in the steady state, (ii) MI induces Ly6C^hi^ monocyte priming towards expression of granulocyte associated genes and transcripts involved in inflammatory activation (*Socs3*), (iii) IFNIC-type monocytes circulate in the steady state and are not induced by MI, (iv) acquisition of monocyte/macrophage states (e.g. *Cxcl3*^*hi*^ monocytes, *Fn1/Ltc4s* macrophages) occurs within the tissue microenvironment rather than during upstream monocyte production and circulation. In particular, our analysis supports the notion that Ly6C^hi^ monocytes follow specific differentiation pathways towards Ly6C^lo^ monocytes in the blood (in line with the idea of Ly6C^lo^ monocytes being terminally differentiated intravascular macrophages (27), (31)), and towards heterogeneous macrophage populations in the ischemic tissue.

### Trajectory inference analysis reveals two main pathways of monocyte to macrophage differentiation in the heart

Having established that cardiac MI-associated macrophage states are not acquired in the periphery, we analyzed monocyte fate once infiltrated in the ischemic heart. To evaluate whether the two major MI-associated macrophage transcriptional states (MI-Mφ-*Trem2/Ifg1* and MI-Mφ-MHCII) result from distinct monocyte differentiation pathways, we performed trajectory inference analysis of monocytes/macrophages in Monocle (32). We used data from our multiplexed analysis of the day 1, 3 and 5 time points (**Figure 2**), as they were the less likely to be biased by batch effect, and showed the best transcript coverage (4,685 detected genes/cell, **Table 1**). Trajectory inference using pseudotemporal ordering requires an assumption on an initial cell state (i.e. the “root” of the trajectory) and direction of the trajectory (33). Here, we based our analysis on the assumption that between day 1 and 5 after MI, macrophages in the infarcted heart essentially derive from recruited Ly6C^hi^ monocytes, a notion supported by previous literature (2),(6) and our own observations. Ordering genes were determined by identifying genes differentially expressed according to the time point of origin of individual cells. Accordingly, cell positions in pseudotime matched their real time appearance in the ischemic heart (**Figure 6A**). Cells corresponding to Ly6C^hi^ monocytes were at the root of the pseudotime tree, while clusters corresponding to Mφ clusters were placed at its end (**Figure 6B**). Mo-*Cxcl3* and Mφ-*Fn1/Ltc4s* preferentially occupied specific early branches of the pseudotime tree, with cells acquiring expression of characteristic genes in function of pseudotime (e.g. *Inhba, Cxcl3, Fn1, Arg1*). At later times, we observed a major trajectory bifurcation (**Figure 6A-C**), leading on one hand to cells highly expressing a set of transcripts including *Trem2, Igf1, Gdf15, Timp2*, or *Ms4a7* (Cell Fate 1, **Figure 6A-D**) while on the other hand cells following the opposite trajectory acquired expression of MHCII encoding genes (Cell Fate 2, **Figure 6A-D**). These results support the notion of two major monocyte-to-macrophage differentiation trajectories in the ischemic heart, giving rise to the *Trem2*^*hi*^*Igf1*^*hi*^ versus MHCII^hi^ gene expression signatures, while a distinct early differentiation pathway gives rise to *Cxcl3*^*hi*^ Mo and *Fn1/Ltc4s* ^*hi*^ Mφ.

**Figure 6:**
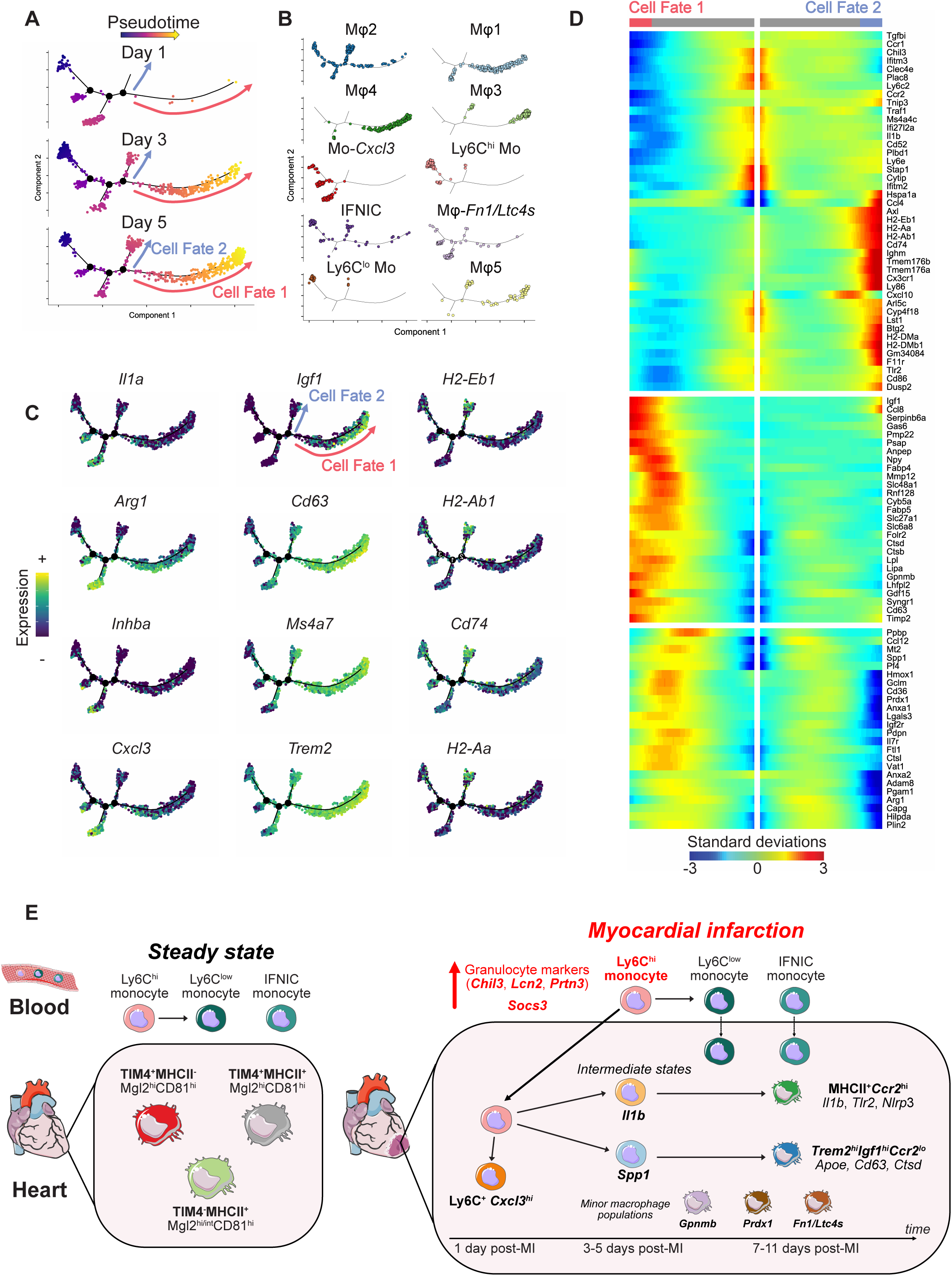
Analysis of monocyte/macrophage differentiation trajectories in the heart. Pseudotime analysis in Monocle split according to **A)** cell time point of origin and **B)** Seurat clusters (see **Figure 2**); **C)** expression of the indicated transcripts projected onto the pseudotime tree; **D)** heatmap showing expression variation of the indicated transcripts (only genes with qval<1.10^−40^ are depicted). **E)** Proposed model of monocyte/macrophage transitions after myocardial infarction.

Altogether, we show that MI induces a shift in circulating Ly6C^hi^ monocytes towards a granulocyte-like state, but that acquisition of tissue-specific signatures of Ly6C^hi^ derived MI-associated macrophages occurs locally. We propose that at 1 day after MI, Ly6C^hi^ monocytes acquire the *Cxcl3*^hi^ state, while at 3 days onwards, they follow two main differentiation pathway giving rise to MHCII^+^*Ccr2*^hi^ and *Trem2*^hi^*Igf1*^hi^*Ccr2*^low^ macrophages. Minor populations of macrophages also accumulate (e.g. Mφ-*Fn1/Ltc4s*), while low numbers of Ly6C^lo^ monocyte and IFNIC monocyte infiltrate the heart over the post-MI time continuum (**Figure 6E**).

## Discussion

Using time-resolved scRNA-seq and CITE-seq analysis of blood and cardiac inflammatory cells in a murine model of MI, as well as by comparing and integrating our results with previous data, we generated a comprehensive single-cell census of cardiac monocyte/macrophage dynamics after MI. We characterized gene expression dynamics during monocyte-to-macrophage transitions and tissue specification in the ischemic heart, uncovered transcriptional regulators that may underlie these processes, and identified new markers that can be used to track monocyte and macrophage subpopulations in the heart.

We recovered cardiac resident macrophages populations, and propose that surface expression of MGL2 and CD81 may help distinguish these cells from recruited monocyte-derived macrophages. Consistent with previous reports, these populations almost entirely vanished from the infarcted heart immediately after MI (2), (5), which was paralleled by infiltration of Ly6C^+^ monocytes. At very early time points after MI, Ly6C^+^ monocytes showed a substantial heterogeneity, with *Cxcl3*^*hi*^ monocytes being preponderant 1 day after MI. This *Cxcl3*^*hi*^ monocyte state appeared to be very transient as these cells could not be observed at substantial levels further than the day 1 time point. Classical Ly6C^hi^ monocytes with a gene expression signature close to blood cells were preponderant thereafter, alongside transition states towards macrophage differentiation. This shows that the acute phase of monocytic inflammation is characterized by the presence of clearly heterogeneous cell populations, rather than a single wave of homogeneous Ly6C^hi^ monocytes (34).

Our data indicates that Ly6C^hi^ monocytes differentiate into two main MI-associated macrophage populations, characterized on one hand by high expression of MHCII encoding genes, and on the other hand by a signature we termed *Trem2*^*hi*^*Igf1*^*hi*^, encompassing other genes potentially involved in resolution of inflammation and ischemic heart repair (e.g. *Timp2, Gdf15, Spp1*) (17), (18), (19). Although we did not directly employ lineage tracing of monocyte-derived versus tissue resident macrophages, several lines of evidence indicate that MI associated *Trem2*^*hi*^*Igf1*^*hi*^ and MHCII^+^ macrophages originate from recruited monocytes. Future experiments with e.g. recently developed *Ms4a3* based lineage tracing of monocytes will allow definitive evaluation of cell origin in the post-MI heart (35). While MI-associated MHCII^+^ macrophages highly expressed *Ccr2, Trem2*^*hi*^*Igf1*^*hi*^ macrophages did not, which may reflect previously described down-regulation of *Ccr2* expression during monocyte to macrophage differentiation (36). Hence, we propose that CCR2 expression cannot be employed to label all differentiated macrophages originating from recruited monocytes. Emergence of other monocyte/macrophage states such as *Cxcl3*^*hi*^ monocytes and *Fn1/Ltc4s*^*hi*^ macrophages may represent a distinct pathway of monocyte differentiation, as these cells shared a number of marker transcripts and activity of specific regulons (e.g. *Cebpb, Hif1a, Klf7*). We furthermore uncovered minor populations characterized by highly specific gene signatures and specific functions, such as *Prdx1* enriched macrophages that may specialize in tissue detoxification and iron handling. Future studies targeting macrophage subsets by e.g. manipulating specific transcriptional regulators, as well as precise localization of cell subsets using spatial transcriptomics methods (37) will uncover the specific functional capacities of these cell populations and their contribution to post-ischemic cardiac repair. Proliferating macrophages were found in the steady state heart, and their proportions drastically dropped at day 1 after MI before reemerging. Analysis of their gene expression signatures suggests that proliferation feeds into all major macrophage subsets. Indeed, proliferating macrophages showed a variety of transcriptional identities indicative of mixed, including monocytic, origins.

Analysis of blood monocytes before and after MI demonstrated that a minute population of IFNIC-type monocytes circulates in the blood already in the steady state, and their proportions did not increase after MI. This is in contrast with a recent report proposing that type I interferon signaling induces remote priming of monocytes in the bone marrow following ischemic injury (30). Besides monocytes, type I interferon signature neutrophils were observed in the steady state blood (38), indicating that several type myeloid cell with the IFNIC signature circulate already in the blood of uninjured mice. The role of type I IFN signaling in post-MI remote priming of monocytes and neutrophils remains to be clarified. We could not observe circulating *Cxcl3*^*hi*^ monocytes or cells with the *Fn1/Ltc4s* signature, indicating that these states are acquired within the ischemic tissue. Direct comparison of blood and heart infiltrating monocytes/macrophages and trajectory reconstruction analysis supported this notion, with tissue-specific acquisition of a set of genes characteristic of MI-associated monocyte/macrophage populations. Ly6C^lo^ monocytes infiltrated the heart without substantial changes in their gene expression profile. Whether their infiltration in the cardiac tissue has any functional consequences remains to be determined. Trajectory reconstruction analysis supported the notion of Ly6C^hi^ to Ly6C^lo^ monocyte conversion as the default fate of circulating monocytes, and that Ly6C^lo^ monocytes do not further differentiate into macrophages in the ischemic heart (10).

Our analysis of circulating monocytes before and after MI indicates that ischemic injury induces a shift towards monocytes enriched for the expression of *Chil3* and several granulocyte-associated genes (e.g. *Prtn3, Lcn2, Wfdc21*). Recent reports proposed that mature monocytes arise from two distinct pathways in the steady state, with monocyte-dendritic progenitors (MDPs) and granulocyte-monocyte progenitors (GMPs) differentiating to monocytes with a “DC-like” state or a “neutrophil-like” state, respectively (29). Our results are thus consistent with a shift towards production of “granulocyte-like” monocytes. This granulocyte like state appears similar to Ym1^+^ Ly6C^hi^ monocytes that emerge after tissue injury (13), as Ym1 is encoded by *Chil3* and Ym1^+^ monocytes are enriched for granulocyte transcripts (*Lcn2, Ngp, S100a8* and *S100a9*) (13). This further corroborates the emergence of a granulocyte-like state in monocytes after tissue injury. Importantly, bulk transcriptome analysis of human monocytes sampled 48 hours after acute MI showed upregulation of *LCN2*, a prototypical marker of the granulocyte-like signature, as well as other granulocyte-associated transcripts (*IL1RN, CXCR1*) (39).

However, our data indicates that analysis of MI-induced emergency myelopoiesis may be affected by surgery-related tissue injury especially at the earliest time points, consistent with previous literature (40). In particular, Iyer et al. (41) showed an approximately 10-fold elevation in the plasma concentration of the emergency myelopoiesis inducing cytokine granulocyte-colony stimulating factor (42) 1 day after sham surgery. Emergence of granulocyte-like monocytes can be induced by LPS (13), (14), indicating that emergence of granulocyte-like monocytes after tissue injury, and in particular cardiac ischemic injury, may reflect damage associated molecular patterns systemically released from the ischemic heart engaging pathways similar to those induced by LPS, i.e. toll-like receptor activation. How this inflammatory priming of monocytes interacts with local cues in the ischemic tissue niche to modulate their downstream differentiation into functionally distinct macrophages populations remains to be determined. In particular, it will be of interest to investigate how innate immune training of myeloid progenitors related to cardiovascular risk factors (e.g. atherogenic conditions (43)) further affect such processes.

Recent reports proposed that pericardial macrophages relocate to the epicardium and have protective properties, inhibiting interstitial fibrosis and preventing cardiac rupture (44), (45). The datasets used here were generated using a classical open-chest MI model. Future studies will be necessary to investigate the transcriptional profile of pericardial macrophages migrating to the heart and how it relates to the populations we described here.

In conclusion, our work provides a novel high-resolution view of the heterogeneity and dynamics of monocyte/macrophage transitions during the acute post-MI inflammation phase, and constitutes a valuable resource for further investigating how these cells may be harnessed and manipulated to promote post-ischemic heart repair.

## Acknowledgements

The work was supported by the Interdisciplinary Center for Clinical Research (IZKF [Interdisziplinäres Zentrum für Klinische Forschung]), University Hospital Würzburg (Project IZKF-E-353 to C.C. and project Z-6 to P.A.), by the German Ministry of Research and Education within the Comprehensive Heart Failure Centre Würzburg (BMBF 01EO1504 to C.C. and A.-E.S.), the Deutsche Forschungsgemeinschaft (DFG, German Research Foundation) (Projektnummer 374031971 – TRR 240 to. A.Z).

## References

1. Frodermann V, Nahrendorf M. 2018. Macrophages and Cardiovascular Health. Physiol Rev 98: 2523–69

2. Dick SA, Macklin JA, Nejat S, Momen A, Clemente-Casares X, Althagafi MG, Chen J, Kantores C, Hosseinzadeh S, Aronoff L, Wong A, Zaman R, Barbu I, Besla R, Lavine KJ, Razani B, Ginhoux F, Husain M, Cybulsky MI, Robbins CS, Epelman S. 2019. Self-renewing resident cardiac macrophages limit adverse remodeling following myocardial infarction. Nat Immunol 20: 29–39

3. Bajpai G, Bredemeyer A, Li W, Zaitsev K, Koenig AL, Lokshina I, Mohan J, Ivey B, Hsiao HM, Weinheimer C, Kovacs A, Epelman S, Artyomov M, Kreisel D, Lavine KJ. 2019. Tissue Resident CCR2- and CCR2+ Cardiac Macrophages Differentially Orchestrate Monocyte Recruitment and Fate Specification Following Myocardial Injury. Circ Res 124: 263–78

4. Chakarov S, Lim HY, Tan L, Lim SY, See P, Lum J, Zhang XM, Foo S, Nakamizo S, Duan K, Kong WT, Gentek R, Balachander A, Carbajo D, Bleriot C, Malleret B, Tam JKC, Baig S, Shabeer M, Toh SES, Schlitzer A, Larbi A, Marichal T, Malissen B, Chen J, Poidinger M, Kabashima K, Bajenoff M, Ng LG, Angeli V, Ginhoux F. 2019. Two distinct interstitial macrophage populations coexist across tissues in specific subtissular niches. Science 363

5. Heidt T, Courties G, Dutta P, Sager HB, Sebas M, Iwamoto Y, Sun Y, Da Silva N, Panizzi P, van der Laan AM, Swirski FK, Weissleder R, Nahrendorf M. 2014. Differential contribution of monocytes to heart macrophages in steady-state and after myocardial infarction. Circ Res 115: 284–95

6. Leuschner F, Rauch PJ, Ueno T, Gorbatov R, Marinelli B, Lee WW, Dutta P, Wei Y, Robbins C, Iwamoto Y, Sena B, Chudnovskiy A, Panizzi P, Keliher E, Higgins JM, Libby P, Moskowitz MA, Pittet MJ, Swirski FK, Weissleder R, Nahrendorf M. 2012. Rapid monocyte kinetics in acute myocardial infarction are sustained by extramedullary monocytopoiesis. J Exp Med 209: 123–37

7. Bajpai G, Schneider C, Wong N, Bredemeyer A, Hulsmans M, Nahrendorf M, Epelman S, Kreisel D, Liu Y, Itoh A, Shankar TS, Selzman CH, Drakos SG, Lavine KJ. 2018. The human heart contains distinct macrophage subsets with divergent origins and functions. Nat Med

8. Dutta P, Sager HB, Stengel KR, Naxerova K, Courties G, Saez B, Silberstein L, Heidt T, Sebas M, Sun Y, Wojtkiewicz G, Feruglio PF, King K, Baker JN, van der Laan AM, Borodovsky A, Fitzgerald K, Hulsmans M, Hoyer F, Iwamoto Y, Vinegoni C, Brown D, Di Carli M, Libby P, Hiebert SW, Scadden DT, Swirski FK, Weissleder R, Nahrendorf M. 2015. Myocardial Infarction Activates CCR2(+) Hematopoietic Stem and Progenitor Cells. Cell Stem Cell 16: 477–87

9. Mouton AJ, DeLeon-Pennell KY, Rivera Gonzalez OJ, Flynn ER, Freeman TC, Saucerman JJ, Garrett MR, Ma Y, Harmancey R, Lindsey ML. 2018. Mapping macrophage polarization over the myocardial infarction time continuum. Basic Res Cardiol 113: 26

10. Hilgendorf I, Gerhardt LM, Tan TC, Winter C, Holderried TA, Chousterman BG, Iwamoto Y, Liao R, Zirlik A, Scherer-Crosbie M, Hedrick CC, Libby P, Nahrendorf M, Weissleder R, Swirski FK. 2014. Ly-6Chigh monocytes depend on Nr4a1 to balance both inflammatory and reparative phases in the infarcted myocardium. Circ Res 114: 1611–22

11. Shiraishi M, Shintani Y, Shintani Y, Ishida H, Saba R, Yamaguchi A, Adachi H, Yashiro K, Suzuki K. 2016. Alternatively activated macrophages determine repair of the infarcted adult murine heart. J Clin Invest 126: 2151–66

12. Satoh T, Nakagawa K, Sugihara F, Kuwahara R, Ashihara M, Yamane F, Minowa Y, Fukushima K, Ebina I, Yoshioka Y, Kumanogoh A, Akira S. 2017. Identification of an atypical monocyte and committed progenitor involved in fibrosis. Nature 541: 96–101

13. Ikeda N, Asano K, Kikuchi K, Uchida Y, Ikegami H, Takagi R, Yotsumoto S, Shibuya T, Makino-Okamura C, Fukuyama H, Watanabe T, Ohmuraya M, Araki K, Nishitai G, Tanaka M. 2018. Emergence of immunoregulatory Ym1(+)Ly6C(hi) monocytes during recovery phase of tissue injury. Sci Immunol 3

14. Yanez A, Coetzee SG, Olsson A, Muench DE, Berman BP, Hazelett DJ, Salomonis N, Grimes HL, Goodridge HS. 2017. Granulocyte-Monocyte Progenitors and Monocyte-Dendritic Cell Progenitors Independently Produce Functionally Distinct Monocytes. Immunity 47: 890–902 e4

15. Stoeckius M, Hafemeister C, Stephenson W, Houck-Loomis B, Chattopadhyay PK, Swerdlow H, Satija R, Smibert P. 2017. Simultaneous epitope and transcriptome measurement in single cells. Nat Methods 14: 865–8

16. Vafadarnejad E, Rizzo G, Krampert L, Arampatzi P, Nugroho VA, Schulz D, Roesch M, Alayrac P, Vilar J, Silvestre J-S, Zernecke A, Saliba A-E, Cochain C. 2019. Time-resolved single-cell transcriptomics uncovers dynamics of cardiac neutrophil diversity in murine myocardial infarction. bioRxiv: 738005

17. Kempf T, Zarbock A, Widera C, Butz S, Stadtmann A, Rossaint J, Bolomini-Vittori M, Korf-Klingebiel M, Napp LC, Hansen B, Kanwischer A, Bavendiek U, Beutel G, Hapke M, Sauer MG, Laudanna C, Hogg N, Vestweber D, Wollert KC. 2011. GDF-15 is an inhibitor of leukocyte integrin activation required for survival after myocardial infarction in mice. Nat Med 17: 581–8

18. Trueblood NA, Xie Z, Communal C, Sam F, Ngoy S, Liaw L, Jenkins AW, Wang J, Sawyer DB, Bing OH, Apstein CS, Colucci WS, Singh K. 2001. Exaggerated left ventricular dilation and reduced collagen deposition after myocardial infarction in mice lacking osteopontin. Circ Res 88: 1080–7

19. Kandalam V, Basu R, Abraham T, Wang X, Soloway PD, Jaworski DM, Oudit GY, Kassiri Z. 2010. TIMP2 deficiency accelerates adverse post-myocardial infarction remodeling because of enhanced MT1-MMP activity despite lack of MMP2 activation. Circ Res 106: 796–808

20. King KR, Aguirre AD, Ye YX, Sun Y, Roh JD, Ng RP, Jr., Kohler RH, Arlauckas SP, Iwamoto Y, Savol A, Sadreyev RI, Kelly M, Fitzgibbons TP, Fitzgerald KA, Mitchison T, Libby P, Nahrendorf M, Weissleder R. 2017. IRF3 and type I interferons fuel a fatal response to myocardial infarction. Nat Med 23: 1481–7

21. Aibar S, Gonzalez-Blas CB, Moerman T, Huynh-Thu VA, Imrichova H, Hulselmans G, Rambow F, Marine JC, Geurts P, Aerts J, van den Oord J, Atak ZK, Wouters J, Aerts S. 2017. SCENIC: single-cell regulatory network inference and clustering. Nat Methods 14: 1083–6

22. Stoeckius M, Zheng S, Houck-Loomis B, Hao S, Yeung BZ, Mauck WM, 3rd, Smibert P, Satija R. 2018. Cell Hashing with barcoded antibodies enables multiplexing and doublet detection for single cell genomics. Genome Biol 19: 224

23. Stuart T, Butler A, Hoffman P, Hafemeister C, Papalexi E, Mauck WM, 3rd, Hao Y, Stoeckius M, Smibert P, Satija R. 2019. Comprehensive Integration of Single-Cell Data. Cell 177: 1888–902 e21

24. Farbehi N, Patrick R, Dorison A, Xaymardan M, Janbandhu V, Wystub-Lis K, Ho JW, Nordon RE, Harvey RP. 2019. Single-cell expression profiling reveals dynamic flux of cardiac stromal, vascular and immune cells in health and injury. Elife 8

25. The Gene Ontology C. 2019. The Gene Ontology Resource: 20 years and still GOing strong. Nucleic Acids Res 47: D330–D8

26. Yu G, Wang LG, Han Y, He QY. 2012. clusterProfiler: an R package for comparing biological themes among gene clusters. OMICS 16: 284–7

27. Mildner A, Schonheit J, Giladi A, David E, Lara-Astiaso D, Lorenzo-Vivas E, Paul F, Chappell-Maor L, Priller J, Leutz A, Amit I, Jung S. 2017. Genomic Characterization of Murine Monocytes Reveals C/EBPbeta Transcription Factor Dependence of Ly6C(-) Cells. Immunity 46: 849–62 e7

28. Gao E, Lei YH, Shang X, Huang ZM, Zuo L, Boucher M, Fan Q, Chuprun JK, Ma XL, Koch WJ. 2010. A novel and efficient model of coronary artery ligation and myocardial infarction in the mouse. Circ Res 107: 1445–53

29. Weinreb C, Rodriguez-Fraticelli A, Camargo FD, Klein AM. 2020. Lineage tracing on transcriptional landscapes links state to fate during differentiation. Science

30. Calcagno DM, Ng RP, Toomu A, Zhang C, Huang K, Aguirre AD, Weissleder R, Daniels LB, Fu Z, King KR. 2020. Type I interferon responses to ischemic injury begin in the bone marrow of mice and humans and depend on Tet2, Nrf2, and Irf3. bioRxiv: 765404

31. Ginhoux F, Jung S. 2014. Monocytes and macrophages: developmental pathways and tissue homeostasis. Nat Rev Immunol 14: 392–404

32. Trapnell C, Cacchiarelli D, Grimsby J, Pokharel P, Li S, Morse M, Lennon NJ, Livak KJ, Mikkelsen TS, Rinn JL. 2014. The dynamics and regulators of cell fate decisions are revealed by pseudotemporal ordering of single cells. Nat Biotechnol 32: 381–6

33. Tritschler S, Buttner M, Fischer DS, Lange M, Bergen V, Lickert H, Theis FJ. 2019. Concepts and limitations for learning developmental trajectories from single cell genomics. Development 146

34. Nahrendorf M, Swirski FK, Aikawa E, Stangenberg L, Wurdinger T, Figueiredo JL, Libby P, Weissleder R, Pittet MJ. 2007. The healing myocardium sequentially mobilizes two monocyte subsets with divergent and complementary functions. J Exp Med 204: 3037–47

35. Liu Z, Gu Y, Chakarov S, Bleriot C, Kwok I, Chen X, Shin A, Huang W, Dress RJ, Dutertre CA, Schlitzer A, Chen J, Ng LG, Wang H, Liu Z, Su B, Ginhoux F. 2019. Fate Mapping via Ms4a3-Expression History Traces Monocyte-Derived Cells. Cell 178: 1509–25 e19

36. Phillips RJ, Lutz M, Premack B. 2005. Differential signaling mechanisms regulate expression of CC chemokine receptor-2 during monocyte maturation. J Inflamm (Lond) 2: 14

37. Rodriques SG, Stickels RR, Goeva A, Martin CA, Murray E, Vanderburg CR, Welch J, Chen LM, Chen F, Macosko EZ. 2019. Slide-seq: A scalable technology for measuring genome-wide expression at high spatial resolution. Science 363: 1463–7

38. Xie X, Shi Q, Wu P, Zhang X, Kambara H, Su J, Yu H, Park S-Y, Guo R, Ren Q, Zhang S, Xu Y, Silberstein LE, Cheng T, Ma F, Li C, Luo HR. 2019. Single-cell transcriptome profiling reveals neutrophil heterogeneity and orchestrated maturation during homeostasis and bacterial infection. bioRxiv: 792200

39. Ruparelia N, Godec J, Lee R, Chai JT, Dall’Armellina E, McAndrew D, Digby JE, Forfar JC, Prendergast BD, Kharbanda RK, Banning AP, Neubauer S, Lygate CA, Channon KM, Haining NW, Choudhury RP. 2015. Acute myocardial infarction activates distinct inflammation and proliferation pathways in circulating monocytes, prior to recruitment, and identified through conserved transcriptional responses in mice and humans. Eur Heart J 36: 1923–34

40. Hoffmann J, Ospelt M, Troidl C, Voss S, Liebetrau C, Kim WK, Rolf A, Wietelmann A, Braun T, Troidl K, Sadayappan S, Barefield D, Hamm C, Nef H, Mollmann H. 2014. Sham surgery and inter-individual heterogeneity are major determinants of monocyte subset kinetics in a mouse model of myocardial infarction. PLoS One 9: e98456

41. Iyer RP, de Castro Bras LE, Cannon PL, Ma Y, DeLeon-Pennell KY, Jung M, Flynn ER, Henry JB, Bratton DR, White JA, Fulton LK, Grady AW, Lindsey ML. 2016. Defining the sham environment for post-myocardial infarction studies in mice. Am J Physiol Heart Circ Physiol 311: H822–36

42. Mitroulis I, Kalafati L, Hajishengallis G, Chavakis T. 2018. Myelopoiesis in the Context of Innate Immunity. J Innate Immun 10: 365–72

43. Christ A, Gunther P, Lauterbach MAR, Duewell P, Biswas D, Pelka K, Scholz CJ, Oosting M, Haendler K, Bassler K, Klee K, Schulte-Schrepping J, Ulas T, Moorlag S, Kumar V, Park MH, Joosten LAB, Groh LA, Riksen NP, Espevik T, Schlitzer A, Li Y, Fitzgerald ML, Netea MG, Schultze JL, Latz E. 2018. Western Diet Triggers NLRP3-Dependent Innate Immune Reprogramming. Cell 172: 162–75 e14

44. Mylonas KJ, Jackson-Jones LH, Andrews JPM, Magalhaes MS, Meloni M, Joshi NV, Allen JE, Newby DE, Dweck MR, Gray GA, Bénézech C. 2019. The pericardium promotes cardiac repair and remodelling post-myocardial infarction. bioRxiv: 771154

45. Deniset JF, Belke D, Lee WY, Jorch SK, Deppermann C, Hassanabad AF, Turnbull JD, Teng G, Rozich I, Hudspeth K, Kanno Y, Brooks SR, Hadjantonakis AK, O’Shea JJ, Weber GF, Fedak PWM, Kubes P. 2019. Gata6(+) Pericardial Cavity Macrophages Relocate to the Injured Heart and Prevent Cardiac Fibrosis. Immunity 51: 131–40 e5

