## Supplementary material for "Single-cell transcriptomic profiling maps monocyte/macrophage transitions after myocardial infarction in mice": Online_Supplement

### Materials and methods

#### ***Data availability***

Data generated for this report have been deposited in Gene Expression Omnibus (GSE135310) and will be made available upon publication. The datasets can browsed in a web-accessible interface:

**Figure 1:** https://infection-atlas.org/4099491356/
**Figure 2:** Monocytes/macrophages alone : https://infection-atlas.org/9074526738/
Monocytes/macrophages and neutrophils: https://infection-atlas.org/4629609923/
**Figure 5:** https://infection-atlas.org/6970782209/

#### ***Animal Models***

Myocardial infarction was induced in 8 to 10 week old male C57BL6/J mice (Janvier) by permanent ligation of the left descendent coronary artery. Mice were exposed to a 12/12 light cycle with lights on at 7:00 and lights off at 19:00. Surgeries were performed between 9:00 and 13:00. Mice received Buprenorphin, 0.1mg/kg i.p. as analgesia, and anesthesia was induced by isoflurane inhalation (4.0%). Mice under deep anesthesia (as revealed by absence of the paw withdrawal reflex) were intubated with an endotracheal cannula and placed under mechanical ventilation (CWE INC. SAR-830-AP; 130 respirations/min, peak pressure max. 18 cm $H_2O$) on a heating pad to maintain body temperature. Anesthesia was maintained with 1.5-2.5% isoflurane during the surgery. The heart was exposed via a 4$^{th}$ intercostal thoracotomy. The left descendent coronary artery was visualized and ligated with a 7/0 non-resorbable nylon suture. The thorax was closed with 3 separated 6/0 non-resorbable nylon sutures and the skin with a continuous 6/0 non-resorbable nylon suture. Buprenorphin was injected i.p. 6 hours after the surgery and twice daily in the two days following surgery. All animal studies and numbers of animals used conform to the Directive 2010/63/EU of the European Parliament and have been approved by the appropriate local authorities (Regierung von Unterfranken, Würzburg, Germany, Akt.-Z. 55.2-DMS-2532-2-743).

#### ***Flow cytometry analysis***

Hearts were enzymatically digested in RPMI medium containing 450U/ml collagenase I (Sigma-Aldrich C0130), 125U/ml collagenase XI (Sigma-Aldrich C7657), 60U/ml Hyaluronidase (Sigma-Aldrich H3506) for 1 hour at 37°C with agitation, washed in PBS/1% FCS and passed through a 70µm cell strainer to generate cardiac cell suspensions. Prior to labeling, cell suspensions were incubated at 4°C for 10 minutes with anti-CD16/32 (Biolegend, Clone 93, 10µg/ml) to block unspecific binding to Fc receptor and then stained at 4°C for 30 minutes with Fixable Viability Dye (ThermoFisher, 1:1000) and fluorochrome/biotin conjugated antibodies against anti-CD45 AlexaFluor700 (BioLegend, Clone 30-F11, 1.2µg/ml), anti-CD11b PercpCy5.5 (BioLegend, Clone M1/70, 0.6µg/ml), anti-Ly6C BV510 (BioLegend, Clone HK1.4, 0.2 µg/ml), anti-Ly6G BrilliantViolet650 (BioLegend, Clone 1A8, 0.6 µg/µl), anti-CD81 Biotin (BioLegend, Clone Eat-2, 5 µg/ml), anti-F4/80 PECy7 (BioLegend, Clone BM8, 2 µg/ml), anti-MHCII BrilliantViolet785 (BioLegend, Clone M5/114.15.2, 0.6 µg/ml), anti-TIMD4 PE (ThermoFisher, Clone 54(RMT4-54), 2 µg/ml), anti-CD301b/MGL2 eFluor660 (ThermoFisher, Clone 11A10-B7, 2µg/ml), Streptavidin BrilliantViolet421 (BioLegend, 0.5 µg/ml). Samples were acquired using a FACS Celesta (BD Bioscience) and analyzed with FlowJo.

***Quantitative Real-Time Polymerase Chain Reaction analysis of bone marrow monocytes***

Total bone marrow cells from one femur per mouse were isolated, FcBlock was performed and cells labeled with Fixable viability dye e780, CD45.2-BV421, CD11b-Percp-Cy5.5, CD115-APC, Ly6G-Alexa488, Ly6C-BV510, SiglecF-PE and Ter119/CD3/CD19/NK1.1-PE-Cy7.

Ly6C$^{hi}$ Monocytes were sorted as singlets/live/CD45$^{+}$/CD11b$^{+}$/(Ter119/CD3/CD19/NK1.1)$^{neg}$/CD115$^{+}$/Ly6G$^{neg}$/Ly6C$^{hi}$. After verifying the sorting purity achieved by our sorting parameters, we sorted 50,000 monocytes directly into the RA1 lysis buffer (with added β-mercaptoethanol) of the Nucleo Spin RNA extraction Kit (Macherey-Nagel, Düren, Germany). Total RNA was then extracted in accordance with the manufacturer's instructions. Equal amounts of template RNA were used for cDNA synthesis, and RNA was reverse transcribed using random hexamer primers of the First Strand cDNA Synthesis kit (ThermoFisher ref. K1612). Quantitative SYBR-green (PowerUp™ SYBR™ Green Master Mix, Applied Biosystems ref. A25742) real-time polymerase chain reaction was performed on triplicate samples of template cDNA on an Applied Biosystems™ QuantStudio™ 6 Flex Real-Time PCR System using specific primer pairs. Quantitative measurements were determined using the ΔCt method, with *Hprt* as the housekeeping gene.

Primers sequences were as followed: *Hprt* forward 5'-TCCTCCTCAGACCGCTTTT-3', reverse 5'-CCTGGTTCATCATCGCTAATC-3'; *Prtn3* forward 5'-CCAGACTTCCGGAGCGATCA-3', reverse 5'-CTGCTGGGGCAGAGAA-3'; *Socs3* forward 5'-ATTTCGCTTCGGGACTAGC-3', reverse 5'-AACTTGCTGTGGGTGACCAT-3'; *Lcn2* forward 5'-CCAGACTTCCGGAGCGATCA-3', reverse 5'-TGTTCTGATCCAGTAGCGACA-3'; *Mmp8* forward 5'- CCATGCCTTCCCAGTACCT-3', reverse 5'-GCACTCCACATCGAGGCATT-3'; *Chil3* forward 5'- ATGGAAGTTTGGACCTGCCC-3', reverse 5'- CCTTGGAATGTCTTTCTCCACAG-3'.

***Single-Cell RNA-seq: sample preparation, sequencing and analysis***

***Dataset in Figure 1:*** *Cell sorting: scRNA-seq kinetics day 0-1-3-5-7*

Methods regarding acquisition of datasets in Figures 1 and 2 have been described in detail in reference (16). Briefly, hearts in the steady state and at various time points after infarction (1, 3, 5 and 7 days) were enzymatically digested in RPMI medium containing 450U/ml collagenase I (Sigma-Aldrich C0130), 125U/ml collagenase XI (Sigma-Aldrich C7657), 60U/ml Hyaluronidase (Sigma-Aldrich H3506) for 1 hour at 37°C with agitation. Viable $CD45^+$ cells were FACS-sorted using a FACS Aria III (BD Biosciences) with a 100µm nozzle, counted and directly loaded into the 10X Genomics Chromium. To ensure that cell preparations did not contain circulating blood leukocytes, mice received an intravenous injection of fluorochrome conjugated anti-CD45.2 antibodies and labeled cells were excluded from sorting.

***Dataset in Figure 2:*** *Cell hashing/CITE-Seq: heart Day 1/3/5 post-myocardial infarction*

Mice received an i.v. injection 5µg of anti-CD45.2-APC (clone 104, ThermoFisher Scientific) in 100µl of PBS under isoflurane anesthesia and were sacrificed by cervical dislocation 5-10 minutes later. Myocardial infarction (at 1 (n=5), 3 (n=5) and 5 days post-MI (n=3)) induction was macroscopically validated and mice were perfused with intracardiac PBS until liver bleaching and the hearts collected. The viable myocardium above the ligation and the right ventricle were removed. The infarcted area, its border zone and the adjacent viable myocardium were processed. The hearts were enzymatically digested as above, and after a Fc-Block step (10µg/ml purified rat anti-mouse CD16/CD32, Biolegend TruStain FcX anti-mouse), the cell suspensions were labeled with Fixable Viability Staining e780 (1:1000) and CD11b (M1/70) antibodies: CD11b-BV510 (Day 1 cells, 1:300, BD Biosciences), CD11B-Percp-Cy5.5 (Day 3 cells, 1:300, BD Biosciences) or CD11b-PE (Day 5 cells, 1:300, BD Biosciences), washed and pooled. Viable cells labeled with CD11b-BV510, CD11b-Percp-Cy5.5 and CD11b-PE were separately sorted using a FACS Aria III (BD Biosciences) with a 100µm nozzle. After sorting, cells were separately labeled with hashtag antibodies (Biolegend TotalSeq-A antibodies, Day 1: Hashtag 1, Day 3: Hashtag 2, Day 5: Hashtag 3) and CITE-Seq antibodies (Biolegend TotalSeq-A against mouse Ly6C, Ly6G, CD64, F4/80, CD11c, MSR1, IA/IE, TIMD4, CX3CR1, and CCR2). Cells were washed, counted and loaded in the 10x Genomics Chromium at a 1,300 cells/µl concentration with the aim of recovering 10,000 total cells using loading volumes recommended by 10X Genomics.

***Dataset in Figure 5:*** *CITE-seq analysis of blood and cardiac CD11b+ cells*

Cells were isolated from blood or ischemic heart of mice in the following conditions: no surgery control (n=3), sham surgery day 1 (n=3), sham surgery day 3 (n=3), myocardial infarction day 1 (n=5), myocardial infarction day 3 (n=5). Venous blood was collected in EDTA containing collection tubes by retroorbital puncture using heparinized hematocrit capillaries and kept on ice with agitation until further processing. Red blood cells were removed by performing erythrocyte lysis in ACK buffer for 5' on ice twice. Ischemic cardiac tissue from mice with 1 and 3 days old infarcts was mechanically dissociated (16) to preserve cell surface epitopes for CITE-seq labeling. Cells were labeled with hashtag antibodies 1 to 7 (TotalSeqA hashtag antibodies, Biolegend, 1:250 dilution), Fixable Viability Staining e780 (1:1000), B220-PE-Cy7 (clone, final concentration 0.6 µg/ml), NK1.1 PE-Cy7 (clone PK136, 2µg/ml), Ter119-PE-Cy7 (clone Ter119, 0.6 µg/ml), CD11b-PercpCy5.5 (M1/70, 0.6µg/ml), Ly6G-PacificBlue (1A8, 1.25µg/ml), SiglecF-Biotin (clone EA798, Miltenyi, 3µg/ml). At this step, CITE-seq antibodies against Ly6G (1.25µg/ml) and CD11b (1.25µg/ml) were also included in the labeling mix to avoid epitopes being inaccessible in the second CITE-seq labeling step. Viable Ter119$^{-}$B220$^{-}$ NK1.1$^{-}$CD11b$^{+}$ cells were sorted using a FACS Aria III (BD Biosciences) with a 100µm nozzle. The final content of Ly6G$^{+}$ neutrophils and Ly6G$^{-}$ non-neutrophils in the sorted cells was adjusted to approximately 1:1 proportions. After sorting, cells were washed once and cells from each sample were pooled together (the volume of each sample used to pool cells was adjusted based on the number of sorted events, to ensure an approximately equal proportion of cells from each sample in the final data). Pooled cells were labeled with TotalSeq-A antibodies (Biolegend) directed against CD64 (5µg/ml), CD54/ICAM1 (5µg/ml), CD49d (5µg/ml), anti-biotin (2.5µg/ml), CD115 (2.5µg/ml), CD62L (1.67µg/ml), Ly6C (1µg/ml), IA-IE (1.25µg/ml), MSR1(5µg/ml), Fcer1a (5µg/ml), CCR3 (5µg/ml) and CXCR4 (5µg/ml), washed twice in PBS/0.04% BSA. Cells were loaded in the 10X Genomics Chromium at a concentration of 1,700 cells/µl with the aim of recovering 15,000 cells.

*scRNA-seq library preparation and sequencing*

Libraries were generated with the Chromium Single Cell 3' Reagents Kit v2 Chemistry (**Dataset in Figure 1**) or v3 Chemistry (**Dataset in Figure 2 and 5**). The v2 chemistry samples were processed following the standard 10x Genomics protocol. The v3 sample was prepared for CITE-Seq/Hashing according to the manual until the cDNA amplification step in which 1 µl (0.2 µM) ADT PCR additive primer to capture Antibody-derived tags (ADTs) and 1 µl (0.1 µM) HTO PCR additive primer to capture Hashtag oligos (HTOs) were added. After cDNA amplification 60 µl (0.6x) SPRI beads (Beckman Coulter) were added to separate the supernatant fraction that contains the ADT/HTO-derived cDNAs (<180bp) and the bead fraction that contains the mRNA-derived cDNAs (>300bp). The mRNA-derived cDNAs (bead fraction) were processed following the standard 10xGenomics protocol. The ADT/HTO-derived cDNAs (supernatant fraction) were purified twice with 2x SPRI. After the two purifications half amount of the eluted cDNAs were amplified for ADTs and half for hashtags. In the same reactions, the ADTs were indexed using TruSeq Small RNA primers and the hashtags using modified TruSeq DNA primers. The libraries were purified once more with 1.6x SPRI (160 µl). Ths method has been developed by Stoeckius et al., 2018 and is described in detail in reference (15) and (22). Additionally, the detailed protocol including the oligo sequences can be accessed here: https://cite-seq.com/protocol. All libraries were quantified by QubitTM 3.0 Fluometer (ThermoFisher) and quality was checked using 2100 Bioanalyzer with High Sensitivity DNA kit (Agilent). Sequencing was performed with S1 or S2 100bp flowcell with Novaseq 6000 platform (Illumina) and the reads for CITE-Seq/Hashing sample were allocated as follows: 5% for the hashtags, 10% for the ADTs and 85% for the mRNAs.

*Single-cell transcriptomics: data analysis*

10X Genomics data, including HTO and ADT libraries, was demultiplexed using Cell Ranger software (version 3.0.1). Mouse GRCm38 reference genome was used for the alignment and counting steps. To evaluate the expression of the cell surface proteins alongside the transcriptome level in our main data set, --feature-ref flag of Cell Ranger software was used which creates a matrix that contains gene expression counts alongside the expression of cell surface proteins. The gene-barcode matrix obtained from Cell Ranger was further analyzed using Seurat v3 (version 2.3.4) (23). For dataset 1, data obtained from each time point was integrated using Cell Ranger. To reduce the batch effect introduced by different sequencing depth, the read depth was equalized between libraries before integrating them.

Datasets in **Figure 2** and **Figure 5** were demultiplexed in Seurat v3 to identify sample of origin of single cells, exclude multiplets (i.e. cells positive for more than one hashtag signal) and cells with no detectable hashtag signal, essentially according to instructions provided by the software developers (https://satijalab.org/seurat/v3.1/hashing_vignette.html). Analysis of cell surface epitope expression by CITE-seq was also performed using a standard Seurat workflow (https://satijalab.org/seurat/v3.1/multimodal_vignette.html). Clustering analysis was performed on RNA levels alone using the standard Seurat clustering workflow (https://satijalab.org/seurat/v3.1/pbmc3k_tutorial.html). Briefly, low quality cells (with more than 5% UMIs mapped to mitochondrial genes) were filtered out, data was log normalized, and 2,000 variable features (i.e. genes with high expression variation across single cells) were identified using the "vst" method. Data was scaled using the "ScaleData" function, and Principal Component Analysis (PCA) was performed. Statistically significant principal components were determined using the JackStraw method. In all datasets, 20 PCs were employed to perform clustering analysis and Uniform Manifold Approximation and Projection (UMAP) dimensional reduction. Immune cell populations (i.e. monocyte/macrophages, dendritic cells, NK cells and neutrophils) were identified based on expression of canonical transcripts and CITE-seq signal of known surface markers (see main text). In the Figure 1, monocyte/macrophages and neutrophils were subset in a new Seurat Object and clustering analysis and UMAP dimension reduction were performed again using 2,000 variable genes and 20 PCs within this data subset. Dataset from **Figure 2** was separated into a “neutrophil” and a “monocyte/macrophage” data subsets, and clustering analysis and UMAP dimension reduction were performed again using 2,000 variable genes and 20 PCs within the respective data subsets. Of note, CITE-seq measurement of some epitopes resulted in poor and uninterpretable signal, notably CCR2 in **Figure 2** and CXCR4 in **Figure 5** dataset. Importantly, we were also unable to obtain satisfactory labelings for flow cytometry analysis of cardiac or circulating immune cells using the same clones from the same supplier (Biolegend), indicating that these clones may not be suitable for CITE-seq analysis as well. CITE-seq measurements of TIMD4, CCR3 or Fcer1a were also poorly conclusive, likely due to expressing cells being mostly absent of the analyzed population. Using the “AddModuleScore” function in Seurat v3, we assigned ISG (Interferon Stimulated Genes) scores using the with the 10 ISGs employed by Calcagno et al. (30) (*Rsad2*, *Ifit2*, *Ifit3*, *Cmpk2*, *Cxcl10*, *Irf7*, *Isg15*, *Oasl1*, *Mx1*, and *Usp18*) as features. To assign “Granulocyte Expression Scores”, the following genes were used as features: *Lcn2*, *Chil3*, *Wfdc21*, *S100a8*, *Wfdc17*, *S100a9*, *Mmp8*, *Elane*, *Retnlg*, *Slpi*, *Ifitm1*, *Grina*.

*Data integration using Canonical Correlation Analysis*

Previously published datasets from Farbehi et al (24) and Dick et al (2) were reanalyzed in Seurat v3. Before performing integration with other datasets, the Farbehi et al (24) data was curated to include only monocyte/macrophages, and “hybrid cells” (expressing both markers of macrophages and non-immune cells) were excluded from analysis. Canonical Correlation Analysis was performed using the “Standard Workflow” from Seurat developers (https://satijalab.org/seurat/v3.1/integration.html) (23). 30 principal components were used to perform clustering and UMAP dimensional reduction.

*Pseudotime analysis in Monocle v2.8*

Differentially expressed genes were determined using the differentialGeneTest function in Monocle (32). For time series data (data in **Figure 6**), differentially expressed genes were determined according to cell time point of origin and 1,071 ordering genes were used. In the blood/heart analysis (**Figure 5**), differentially expressed genes were determined according to cell clusters and 1,076 ordering genes were used. Dimension reduction was performed using the “reduceDimension” function and the “DDRTree” method and pseudotime ordering was performed using the “orderCells” function.

*SCENIC analysis*

SCENIC was applied as described in (21). For the analysis, mm9 motif ranking database supplied in SCENIC package was used to score potential regulons. The input matrix genes that were not present in motif ranking database were excluded from further analysis. Target genes that did not show a positive correlation based on the GENIE3 co-expression algorithm (> 0.03) in each module were filtered out. TF-modules having less than 20 genes were filtered out and the remaining TF-modules were further examined.

**Supplementary text (description of figure S5):**

We reanalyzed recently published scRNA-seq datasets of cardiac macrophages in the steady state heart and at 11 days after infarction (2) (**Figure S5A-C**), as well as non-cardiomyocyte cells in the steady state and at 3 and 7 days after infarction (with a focus on the myeloid cell data subset) (24) (**Figure S5D-F**). We reanalyzed both datasets in Seurat (23), and found cell populations with gene expression patterns clearly reminiscent of our own analyses. In particular, we observed cells corresponding to MHCII$^{+}$ and *Lyve1*$^{+}$ resident macrophages, as well as proliferating macrophages. Post-MI monocyte/macrophages included Ly6C$^{hi}$ (Dick et al. cluster 13, Farbehi et al. cluster 2) and Ly6C$^{low}$ monocytes (Dick et al. cluster 4, Farbehi et al. cluster 9), IFNICs (Dick et al. cluster 12, Farbehi et al. cluster 7), *Fn1/Ltc4s*$^{hi}$ macrophages (Dick et al. cluster 4, Farbehi et al. cluster 10), macrophages enriched for MHCII encoding genes, and cells enriched for a number of genes also expressed in post-MI macrophages in our datasets (e.g. *Apoe*, *Ctsd*, *Trem2*, *Gpnmb*, *Syngr1*, *Fabp5*, *Cd63*, *Cd9*, *Igf1*, *Gpr137b*, *Ctsb*, *Ms4a7;* Dick et al. cluster 0 and 2, Farbehi et al. cluster 1 and 3). B cell and non-leukocyte contamination were observed, in addition to cells enriched for markers of both Mφ and non-immune cells (*Kdr*, *Esam*, *Cdh5;* Farbehi et al. cluster 4) in the Farbehi et al. dataset (24). Although these cells have been proposed as potential “hybrid” Mφ/endothelial cells (24), we failed to detect them in other datasets.

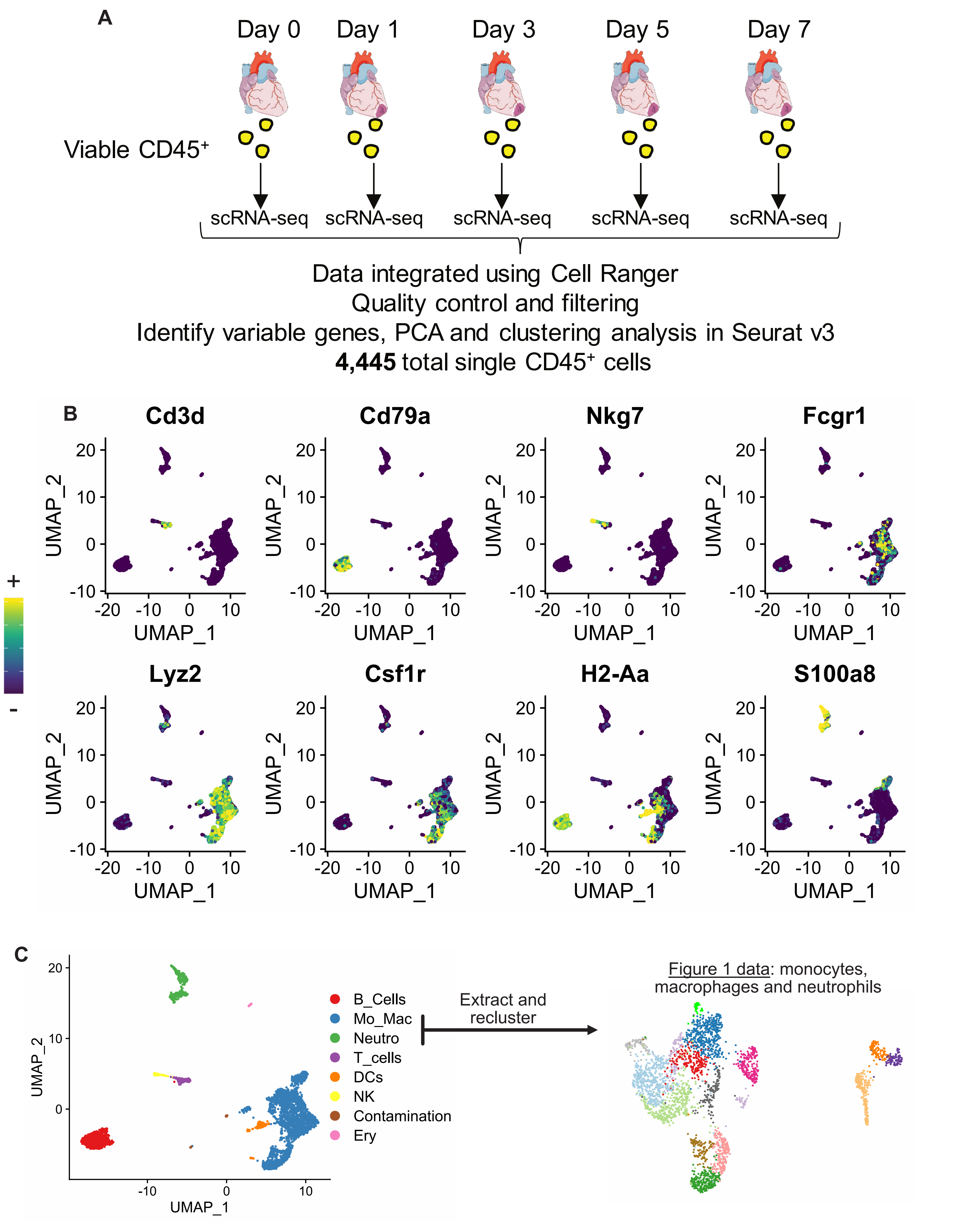


**Figure S1: Experimental design and identification of major immune lineages in total CD45⁺ cells (related to Figure 1). A)** Summary of the experimental design; **B)** expression of transcripts for selected marker genes of major immune lineage projected onto the UMAP plot for total CD45+ cells; **C)** lineage annotation projected onto the UMAP plot. Neutro= neutrophils, DCs=dendritic cells (identified by expression of e.g. *Itgax*, *Cd74*, *Cd209a*, *Flt3*), Mo_Mac=monocytes/macrophages, NK: natural killer cells, Ery=erythrocytes.

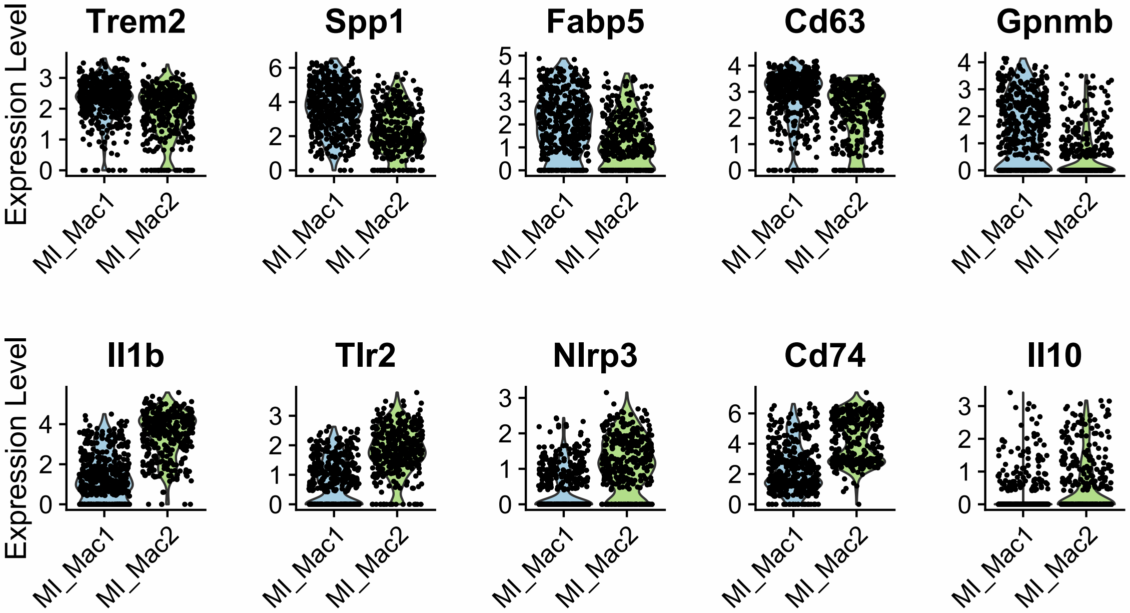


**Figure S2: Gene expression distribution in MI-associated macrophages (related to Figure 1).** Violin plots showing expression level of the indicated transcripts in the two major MI-associated macrophage clusters.

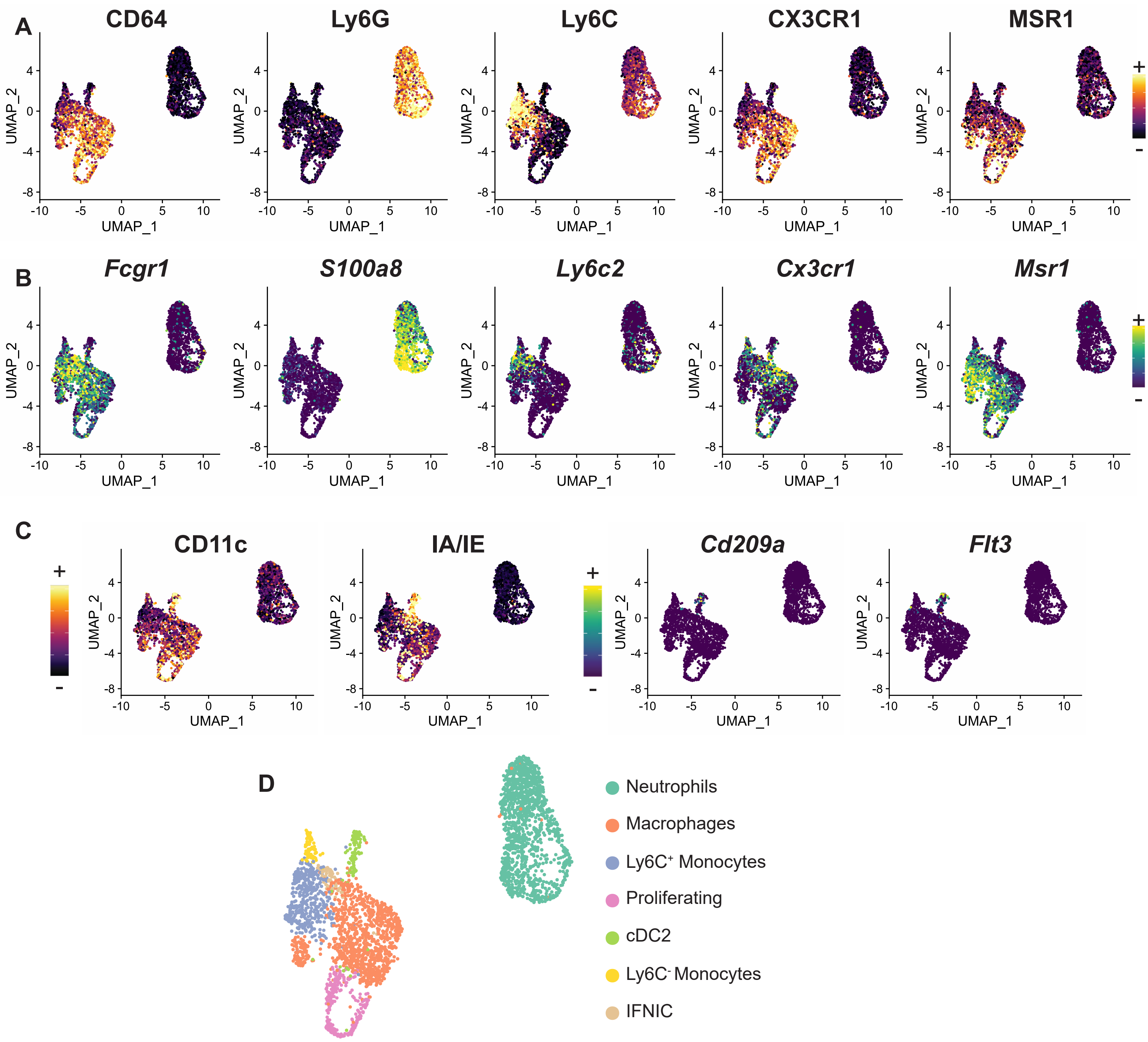


**Figure S3: Multiplexed analysis of cardiac CD11b+ cells at day 1, 3 and 5 after MI (related to Figure 2).** Identification of cells corresponding to neutrophils and monocytes/macrophages based on **A)** cell surface epitope (CITE-seq) signals and **B)** marker gene transcript levels, shown projected on the UMAP plot representation of the dataset; **C)** identification of cDC2 based on expression of surface CD11c and MHCII (IA/IE), and transcript levels of *Cd209a* and *Flt3*; **D)** identification of major cellular identities projected onto the UMAP plot (cDC2: type 2 classical dendritic cells).

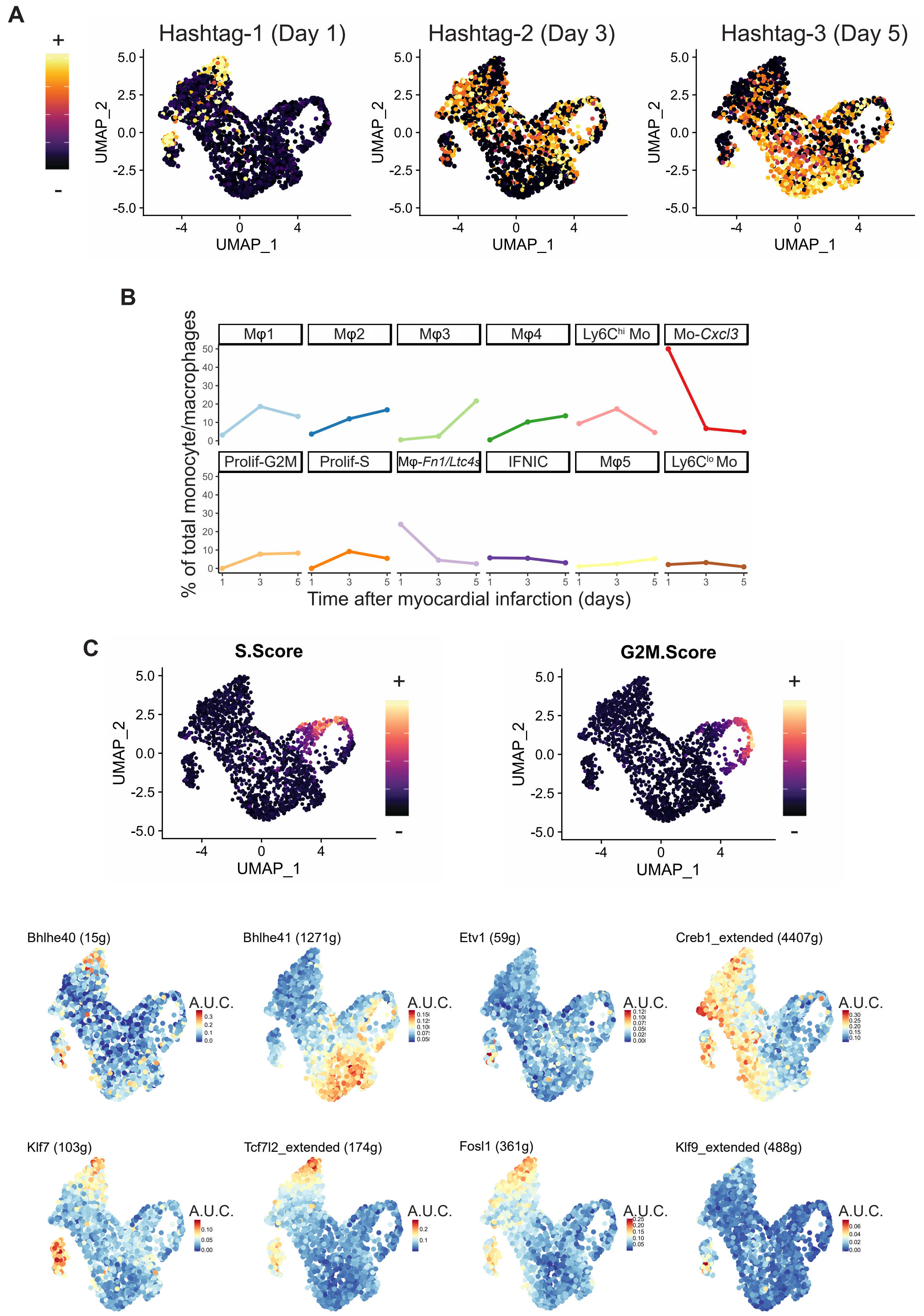


**Figure S4: Multiplexed analysis of monocytes/macrophages at day 1, 3 and 5 after MI (related to Figure 2). A)** Hashtag antibody signal was used to identify cell type point of origin (related to **Figure 2B-C**) and is here projected onto the UMAP plot of the monocyte/macrophage data subset; **B)** proportion of each cluster according to time after MI; **C)** cell cycle phase scoring was performed in Seurat, and scores for S phase and G2M phase are projected onto the UMAP plot of the monocyte/macrophage subset; **D)** activity of the indicated regulons as predicted in SCENIC (A.U.C.: area under the curve) projected onto the UMAP plot. The number of genes associated to each regulon is indicated in brackets.

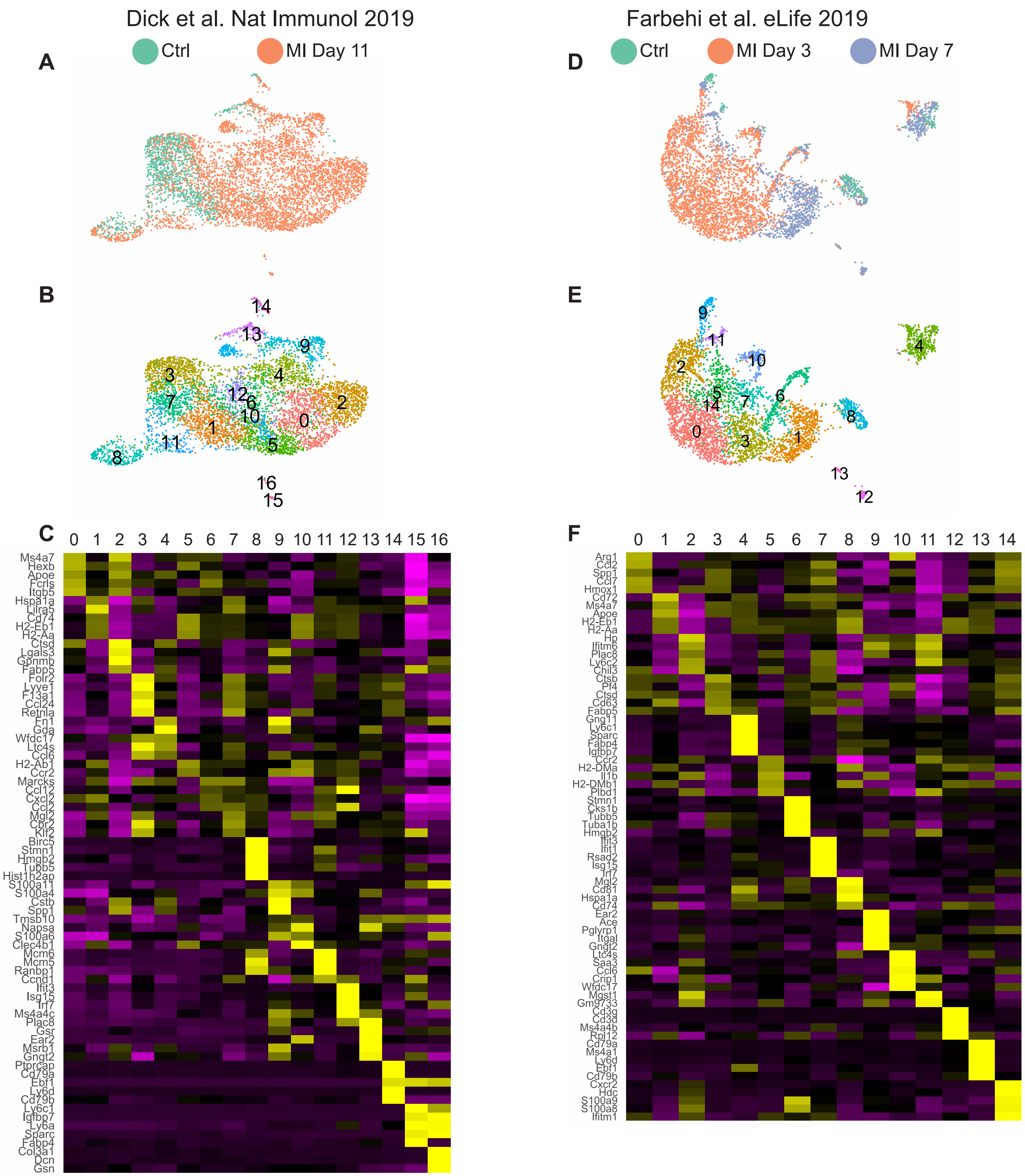


**Figure S5: Reanalysis of scRNA-seq data from Dick et al. Nat Immunol 2019 (A-C, left) and Farbehi et al. eLife 2019 (D-F, right) in Seurat v3 (related to Figure 3). A and D:** time point of origin of single cells projected onto the UMAP plot, **B and E: c**lustering analysis projected onto the UMAP plot, **C and F:** averaged expression heatmap of the top 5 marker genes for each cluster (overlapping markers are shown only once)

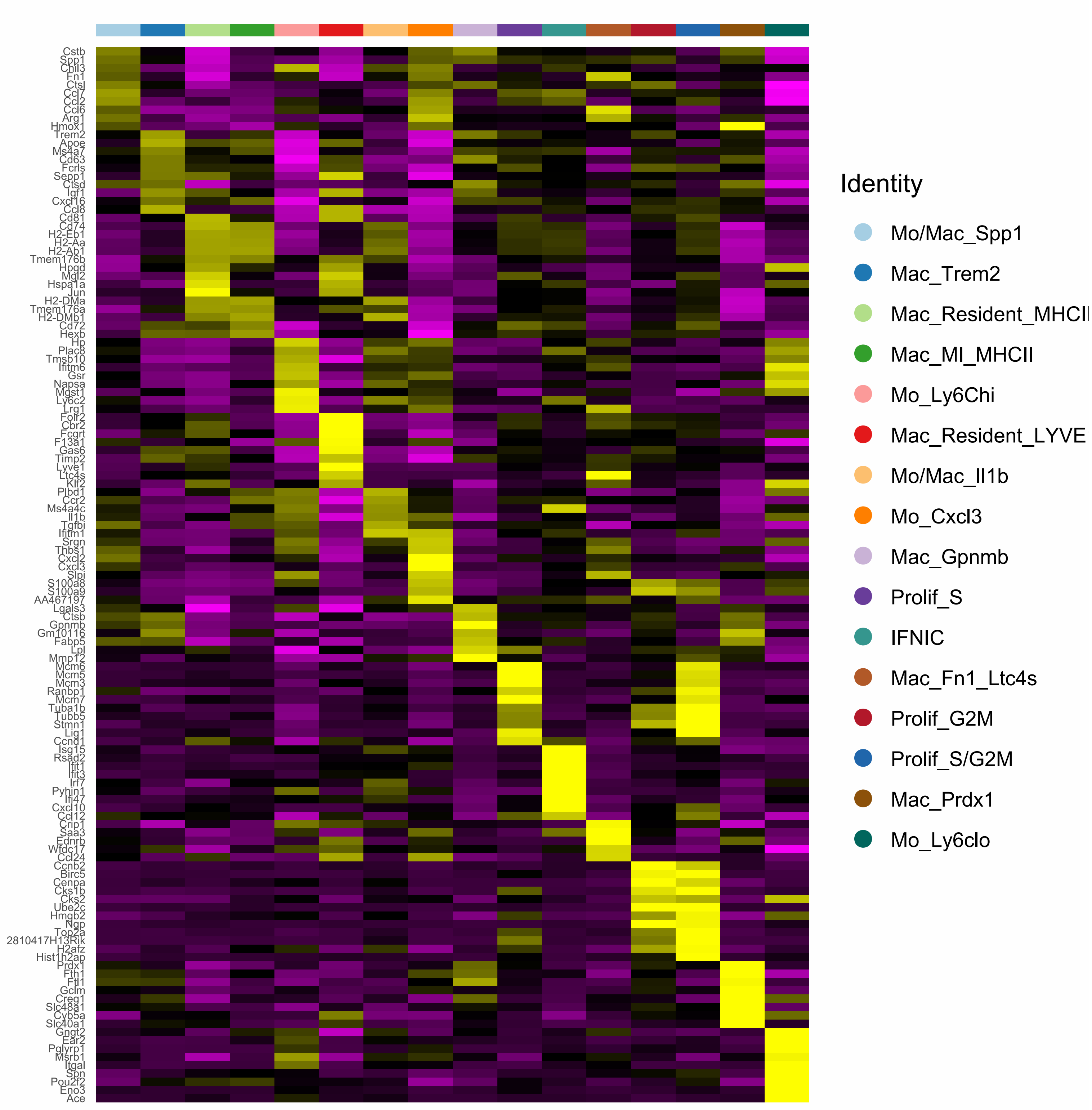


**Figure S6: Gene expression profile of clusters in the CCA-integrated data (related to Figure 3).** Heatmap showing the top 10 enriched genes (ranked by log2 fold change) in each cluster of the CCA-integrated data (average scaled expression). Marker genes overlapping several clusters are listed only once.

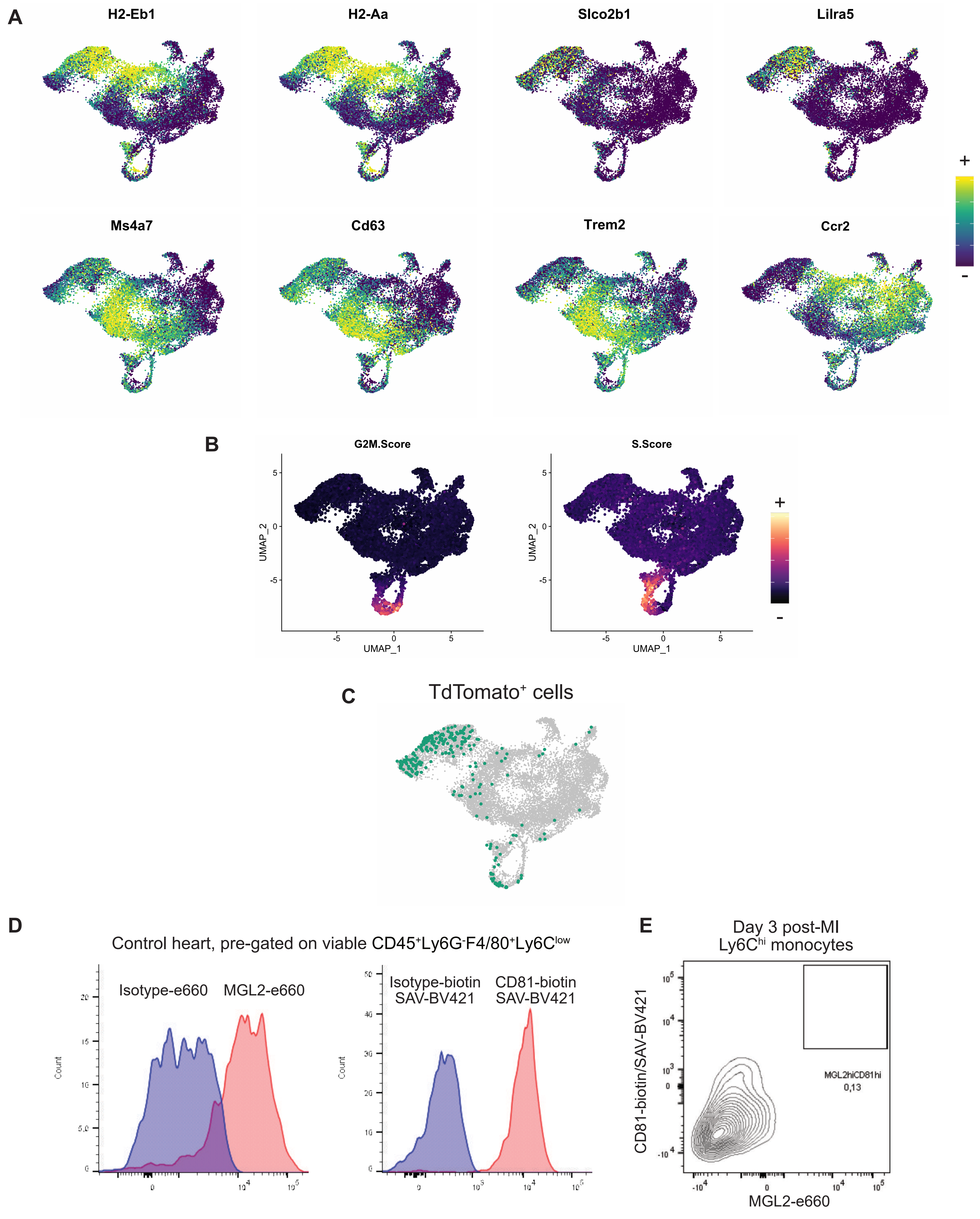


**Figure S7: Cell cycle scoring and gene expression patterns in integrated monocyte/macrophage datasets, day 0 to 11 post-MI (related to Figure 3). A)** Cell cycle scoring for S and G2M phase projected onto the UMAP plot; **B)** Expression of the indicated transcripts projected onto the UMAP plot (please note that for clarity, contrast has been enhanced by applying minimum and maximum expression cutoffs). **C)** cells positive for TdTomato transcripts (from Dick et al. 2019) are highlighted on the UMAP plot; **D)** isotype control validation of MGL2 and CD81 labelings in resident cardiac macrophages (SAV=streptavidin) and **E)** expression of MGL2 and CD81 on Ly6C$^{hi}$ monocytes in the heart at day 3 after infarction.

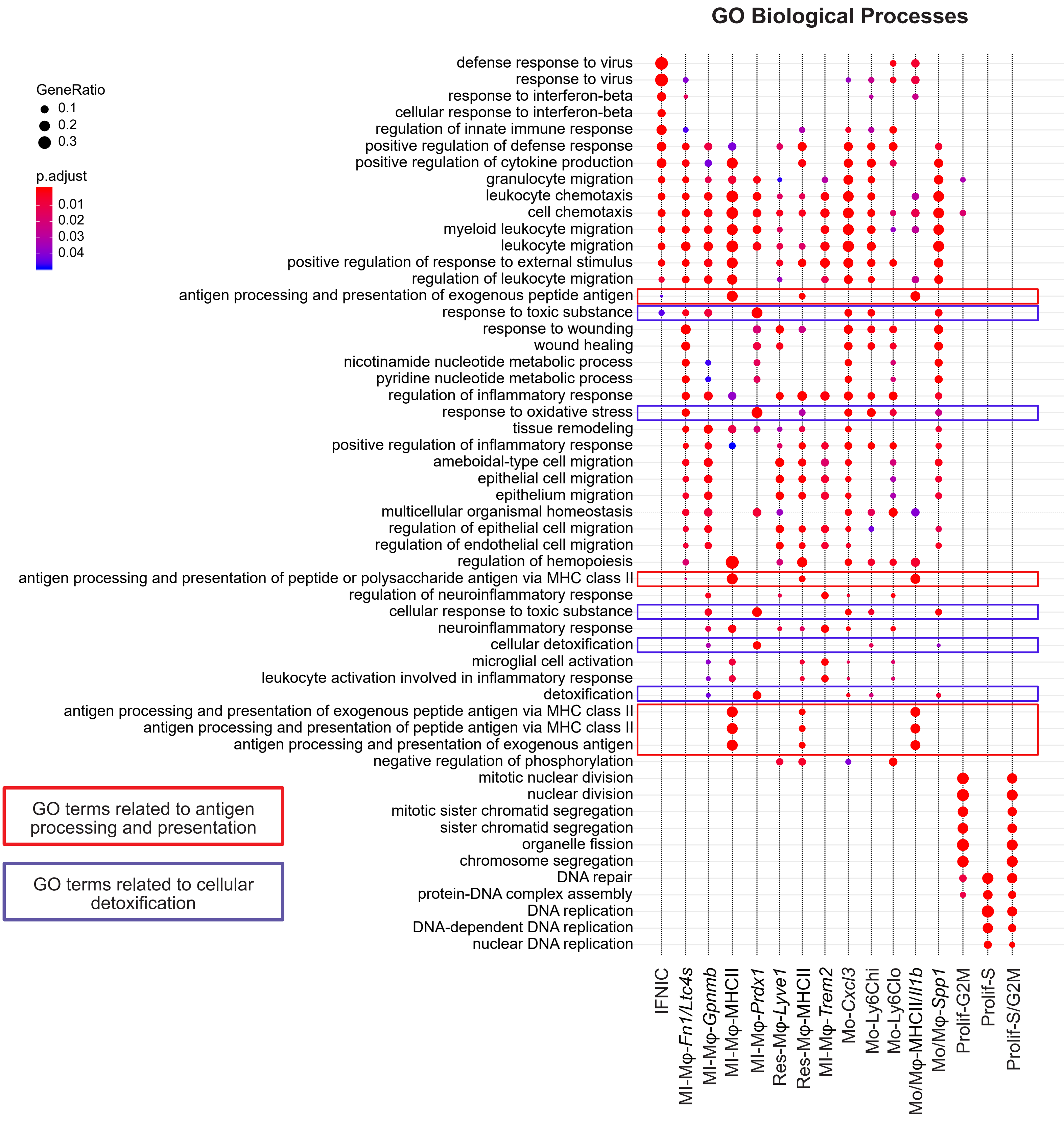


**Figure S8: Gene ontology enrichment analysis for Biological Processes in monocyte/macrophage populations over the post-MI time continuum.** Graphical representation of the most enriched term for Gene Ontology Enrichment analysis-Biological Processes in the 16 clusters found in the CCA-integrated dataset (see Figure 3). The full list of enriched GO-Biological Processes terms can be found in the supplementary Table S4.

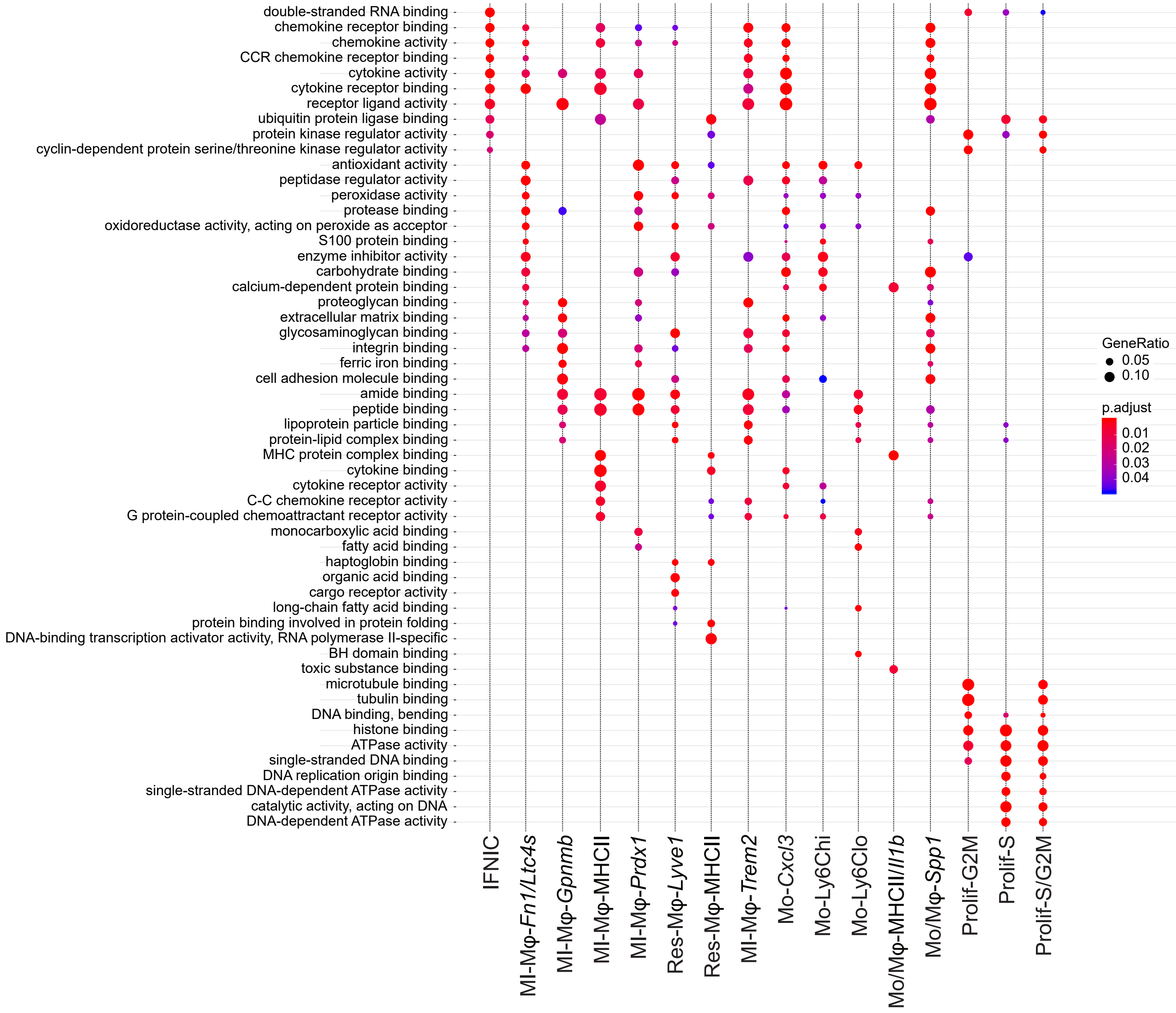


**Figure S9: Gene ontology enrichment analysis for Molecular Function in monocyte/macrophage populations over the post-MI time continuum (related to figure 3).** Graphical representation of the most enriched term for Gene Ontology Enrichment analysis-Molecular Function in the 16 clusters found in the CCA-integrated dataset (see Figure 3). The full list of enriched GO-Molecular Function terms can be found in Table S5.

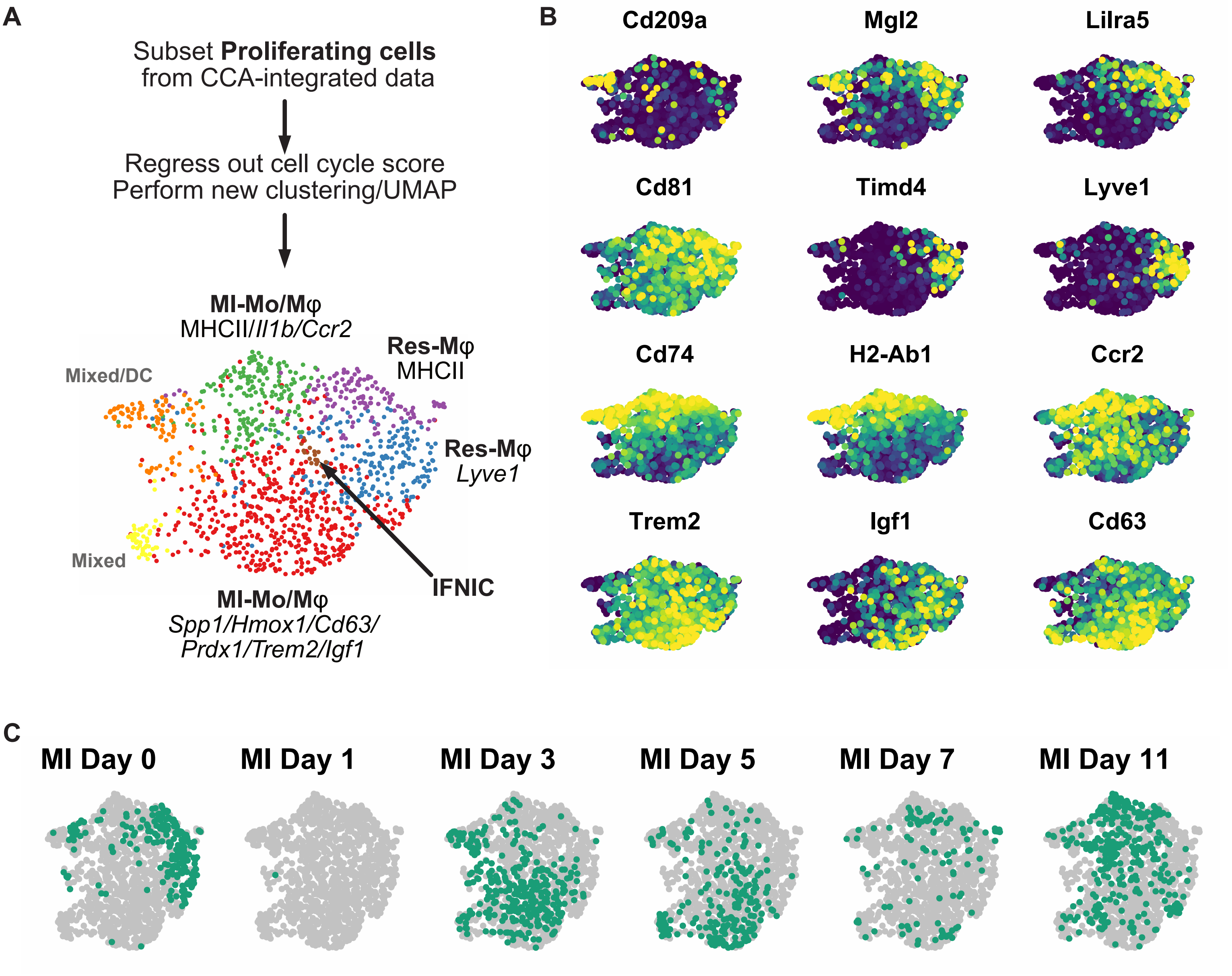


**Figure S10: Identification of proliferating macrophages. A)** proliferating cells were extracted from the integrated data (Figure 3) and cell cycle scoring was regressed out before re-computing clustering analysis and UMAP dimensional reduction, bottom: resulting UMAP and clusters; **B)** expression of the indicated transcripts projected onto the UMAP plot, **C)** Time point of origin of the single cells projected onto the UMAP plot

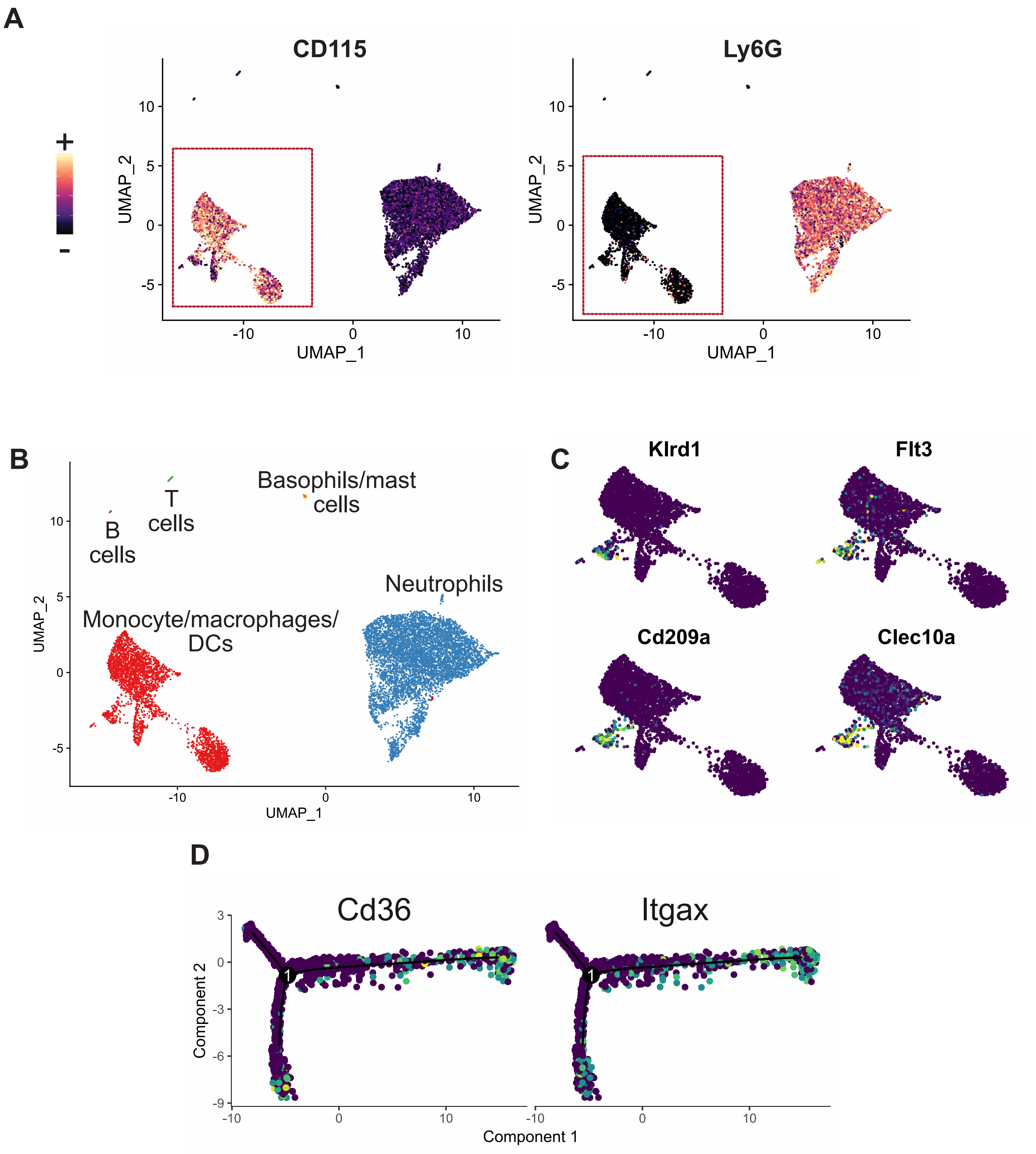


**Figure S11: CITE-seq analysis of blood CD11b+ cells and monocytes (related to Figure 5).** UMAP of single-cell RNA-seq data performed in total viable B220-NK1.1-Ter119-CD11b+ cells sorted from the blood and infarcted heart with **A)** projection of the CITE-seq signal for CD115 and Ly6G (red dashed rectangle: cells corresponding to monocyte/macrophage/DCs) and **B)** identification of the major lineages (including minute contamination by B cells, T cells and mast cells/basophils). **C)** Expression of the indicated cDC2 characteristic transcripts projected onto the UMAP plot in the monocyte/macrophage/DC subset; **D)** expression of the indicated transcripts projected onto the Monocle pseudotime tree (related to **Figure 5G**).

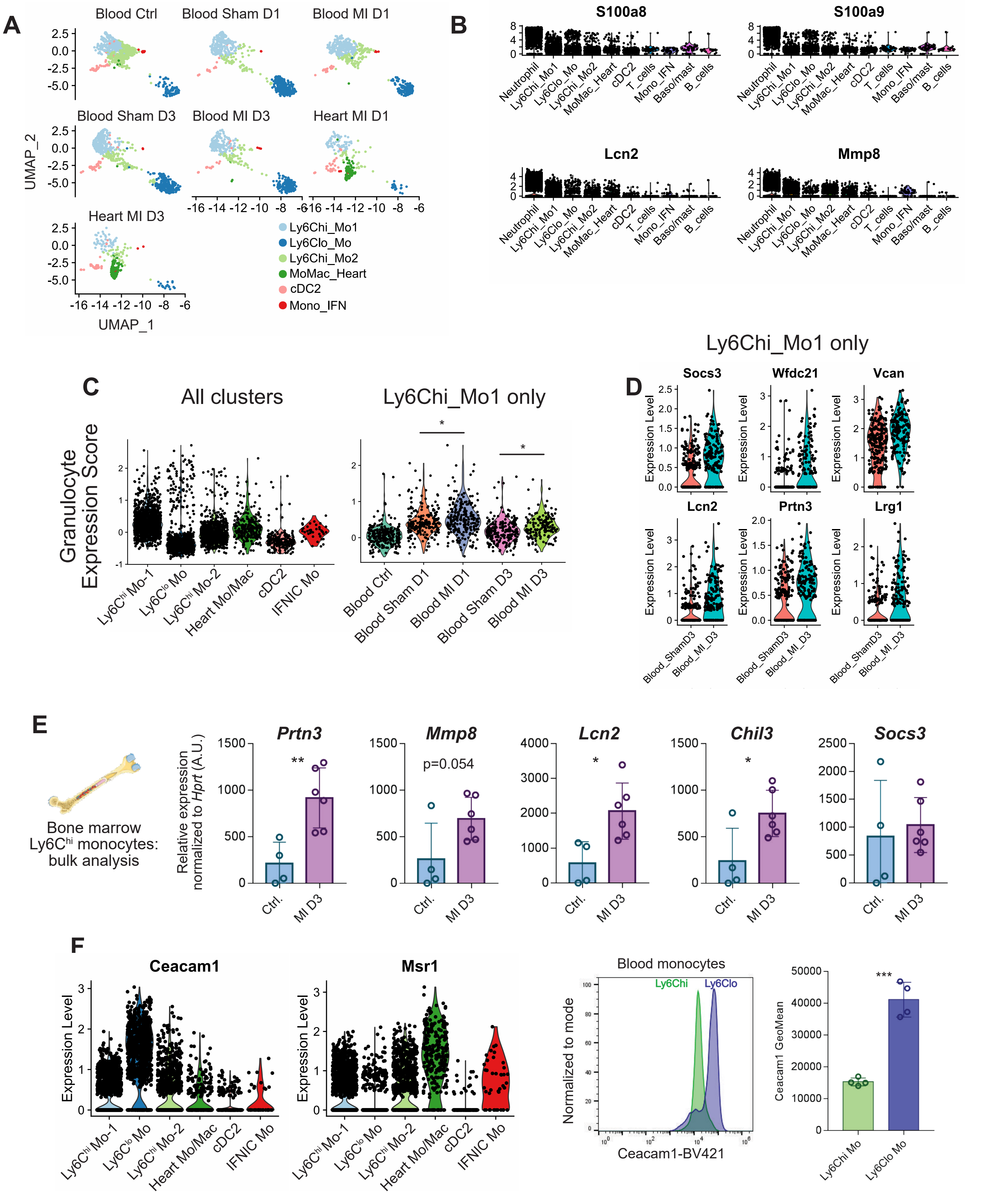


**Figure S12: MI and injury induced priming in blood monocytes (related to Figure 5). A)** UMAP plot split by sample of origin; **B)** violin plots showing expression of characteristic granulocyte transcripts in the indicated cell populations; **C)** violin plot showing the "Granulocyte Expression Score" in monocyte/macrophage/cDC2 clusters in the blood and heart (left) and in blood Ly6Chi_Mo1 cluster according to sample of origin (right; * adjusted p value<0.05); **D)** violin plots showing expression of the indicated transcripts in Ly6Chi_Mo1 cluster in cells from the Blood-Sham-Day 3 and Blood-MI-Day 3 conditions; **E)** expression of the indicated transcripts as measured by qPCR in sorted bone marrow Ly6C$^{hi}$ monocytes (A.U.=arbitrary units); **F)** violin plot showing expression of Ceacam1 and Msr1 in the monocyte/macrophage populations and flow cytometry analysis of Ceacam1 surface level in blood Ly6C$^{hi}$ and Ly6C$^{lo}$ monocytes in the steady state, ***p<0.001 (unpaired t test).

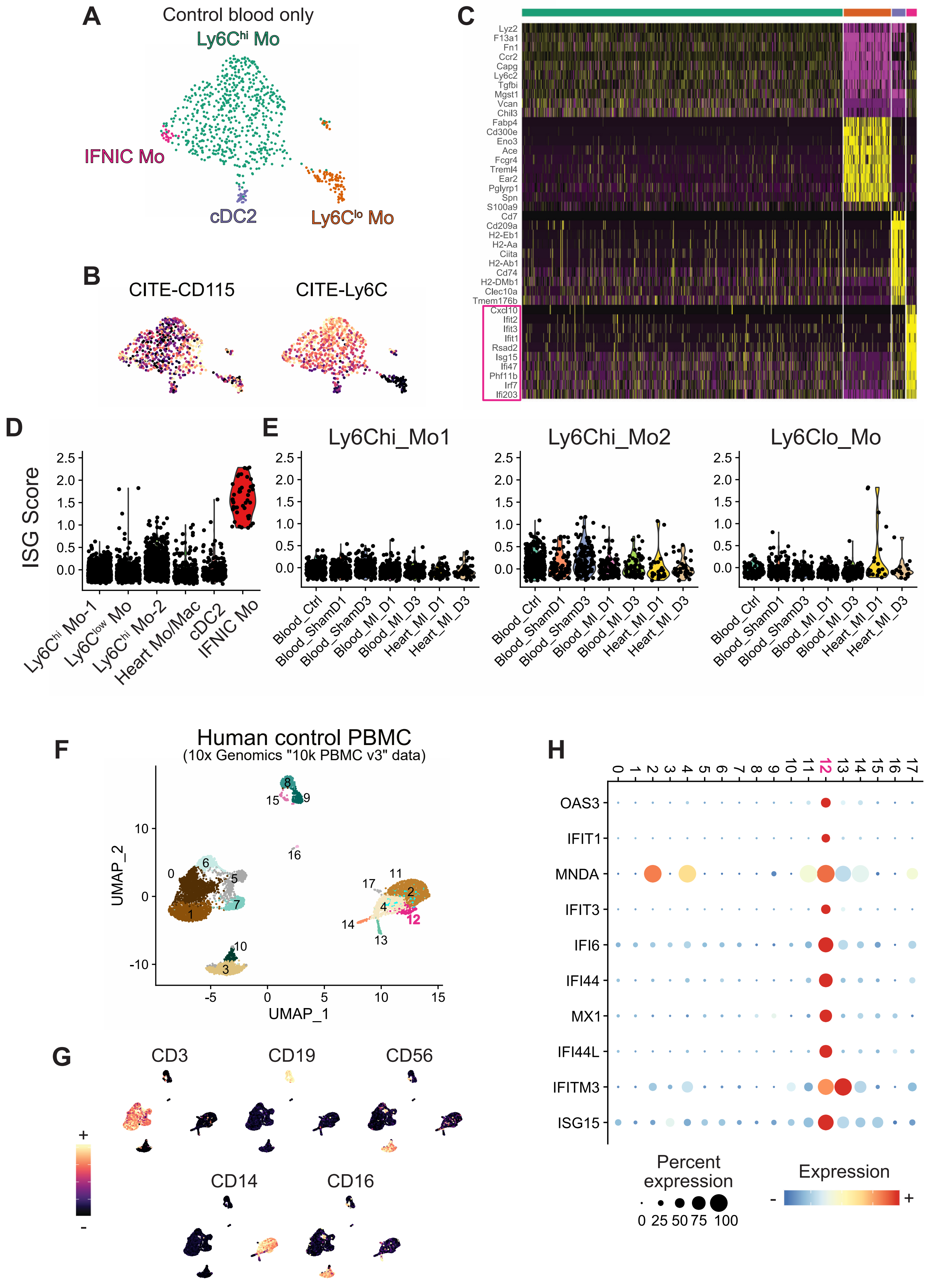


**Figure S13: Type I IFN response monocytes circulate in the steady state**. **A-D**: Data corresponding to blood monocytes and cDC2 from the non-operated control condition (as identified via cell hashing, see Figure 4) were reanalyzed separately in a new unbiased clustering analysis performed at low resolution in Seurat v3 (0.2). **A)** UMAP plot of 723 cells, **B)** CITE-seq signal for CD115 and Ly6C, **C)** heatmap of the top 10 marker genes in each cluster. **D-E)** Type I IFN response score in **D)** all populations from the data shown in Figure 5 and **E)** in monocyte clusters according to sample of origin. **F)** UMAP plot of 10k human PBMCs from a healthy donor (10x Genomics data), with Seurat clustering analysis; **G)** CITE-seq signal for the indicated surface marker projected onto the UMAP plot; **H)** DotPlot showing expression of the indicated transcripts in the PBMC clusters.
